# Modelling spatiotemporal trends in the frequency of genetic mutations conferring insecticide target-site resistance in African malaria vector species

**DOI:** 10.1101/2021.09.15.460499

**Authors:** Penelope A. Hancock, Amy Lynd, Antoinette Wiebe, Maria Devine, Johnathan Essandoh, Francis Wat’senga, Emile Z. Manzambi, Fiacre Agossa, Martin J. Donnelly, David Weetman, Catherine L. Moyes

## Abstract

Resistance in malaria vectors to pyrethroids, the most widely used class of insecticides for malaria vector control, threatens the continued efficacy of vector control tools. Target-site resistance is an important genetic resistance mechanism caused by mutations in the voltage-gated sodium channel (*Vgsc*) gene that encodes the pyrethroid target-site. Understanding the geographic distribution of target-site resistance, and temporal trends across different vector species, can inform strategic deployment of vector control tools. Here we develop a Bayesian statistical spatiotemporal model to interpret species-specific trends in the frequency of the most common resistance mutations, *Vgsc*-995S and *Vgsc*-995F, in three major malaria vector species *Anopheles gambiae, An. coluzzii*, and *An. arabiensis*. For nine selected countries, we develop annual predictive maps which reveal geographically-structured patterns of spread of each mutation at regional and continental scales. The results show associations, as well as stark differences, in spread dynamics of the two mutations across the three vector species. The coverage of ITNs was an influential predictor of *Vgsc* allele frequencies in our models. Our mapped *Vgsc* allele frequencies are a significant partial predictor of phenotypic resistance to the pyrethroid deltamethrin in *An. gambiae* complex populations, highlighting the importance of molecular surveillance of resistance mechanisms.

## INTRODUCTION

A major challenge in malaria control involves managing the threat that insecticide resistance in mosquitoes poses to the efficacy of vector control technologies. Insecticide-based vector control techniques, including indoor residual spraying (IRS) and insecticide-treated bednets (ITNs), are pivotal to malaria prevention, with ITNs in particular being responsible for a large portion of the reductions in malaria cases achieved over the period 2000-2015 (Bhatt *et al*. 2015a). ITNs rely on pyrethroid insecticides, which are used in the treatment of all ITNs pre-approved by the WHO and in many indoor residual sprays still used today (Tangena *et al*. 2019; World Health Organization 2020; Moyes *et al*. 2021). Pyrethroid resistance in malaria vectors has spread extensively throughout Sub-Saharan Africa (Hancock *et al*. 2020), and in 2017 the mosquito sample mortality following exposure to a pyrethroid as measured by a WHO standard susceptibility test had dropped to less than 50% in at least 34 malaria endemic countries.

The prevalence of insecticide resistance phenotypes in African malaria vector species is highly heterogeneous across geographic space (Hancock *et al*. 2020), and underpinned by variation in genetic resistance mechanisms (Miles *et al*. 2017b), which have the potential for rapid long range spread (Clarkson *et al*. 2021). Geographically comprehensive insecticide resistance monitoring and surveillance is therefore essential to track changes in resistance, interpret trends and anticipate upcoming threats. Unfortunately, despite the recommendations of the WHO Global Plan for Insecticide Resistance Management (GPRIM) (World Health Organization 2012) for the instigation of comprehensive and routine insecticide resistance monitoring, the available surveillance data is sparse throughout Sub-Saharan Africa, with 89% of administrative districts having no recorded measurements in the period 2015-2017 (Moyes *et al*. 2020). Standard susceptibility bioassays to measure phenotypic resistance are labour intensive and difficult to scale up. Moreover, where morphologically-cryptic vectors are present, susceptibility bioassays are rarely used to measure resistance at the level of individual species, and do not provide information about mechanisms of resistance. Results can also be sensitive to environmental testing conditions, which are often difficult to standardise in the field (Ismail *et al*. 2018; Weetman *et al*. 2018). Genetic, and in due course genomic, surveillance to track the frequency of variants that are associated with phenotypic resistance is more scalable, insensitive to collection and environmental conditions, and can distinguish between different resistance mechanisms across different vector species.

A major challenge for genetic surveillance lies in identifying variants, or genomic regions, that are important determinants of different types of phenotypic resistance (Donnelly *et al*. 2016). Target-site resistance is an important pyrethroid resistance mechanism in *Anopheles gambiae* complex mosquitoes (Miles *et al*. 2017b; Clarkson *et al*. 2021), and is the most widely monitored genetic mechanism in field malaria vector populations. It is caused by mutations within the *Vgsc* gene that encodes the voltage-gated sodium channel, which is the physiological target of pyrethroid insecticides. Three single point mutations (SNPs) within the *Vgsc* gene are known to confer pyrethroid resistance; these include two substitutions on the 995 codon, L995F (originally named L1014F;Martinez-Torres *et al*. 1998) and L995S (originally named L1014S; Ranson *et al*. 2000), and a third substitution N1570Y (originally named N1575Y; Jones *et al*. 2012). The L995F and L995S mutations occur in the same codon and they cannot co-occur on a single chromosome, while the N1570Y mutation occurs in a different codon and has been found to increase resistance in association with L995F (Jones *et al*. 2012). Genome sequencing has recently identified numerous other non-synonymous SNPs within the *Vgsc* gene, some apparently subject to recent positive selection, indicating that target-site resistance has a complex molecular basis, likely increasingly so over time (Clarkson *et al*. 2021).

The extent to which phenotypic resistance in field malaria vector populations depends on these multifaceted genetic mechanisms remains uncertain (Donnelly *et al*. 2016). Genotype-phenotype association studies are complicated by the polygenic nature of insecticide resistance and the complex population structure of African *Anopheles gambiae* mosquitoes (Clarkson *et al*. 2014; Miles *et al*. 2017a). The *Anopheles gambiae* complex is made up of at least eight individual vector species, five of which are major malaria vectors: *An. gambiae, An. coluzzii, An. arabiensis, An. melus*, and *An. merus* (Wiebe *et al*. 2017; Barron *et al*. 2019; Charlwood 2019). The distribution of the different vector species is geographically heterogeneous, with gradients in species composition occurring across regional and continental scales (Sinka *et al*. 2016). Mechanisms of insecticide resistance differ across these three species (Ranson *et al*. 2011). The evolutionary trajectories of resistance depend on the specific ecology of individual species, the selection pressures present in the environment, and patterns of dispersal, migration and introgression across different populations (Simard *et al*. 2009; Clarkson *et al*. 2014; Fontaine *et al*. 2015; Pombi *et al*. 2017).

Spatial modelling analysis is required to interpret spatial and temporal trends in insecticide resistance surveillance data that monitor the prevalence of different types of resistance in vector species (Hancock *et al*. 2020). This is because sampling locations are heterogeneously distributed across Africa, and variable across sampling times and across the different types of resistance phenotypes and/or genetic mechanisms that were tested in the sample. Geospatial models can quantify geographically explicit temporal trends in resistance (Hancock *et al*. 2020). The ability of geospatial models to extrapolate predictions across unsampled locations can help compensate for sparsity in surveillance data, and allow anticipation of contemporary resistance levels before new surveillance results become available (Moyes *et al*. 2020). Further, geospatial models offer a flexible framework for combining different datasets that describe separate but related aspects of resistance. They can incorporate measures of resistance across different vector species, as well as genetic and phenotypic measures of resistance, within the same modelling framework (Hancock *et al*. 2018). The ability of spatial models to predict resistance can benefit greatly from incorporating information about environmental characteristics such as climate, vegetation and land use; importantly, variables describing the distribution of insecticide-based vector control interventions across the landscape can be included as potential predictors (Hancock *et al*. 2020).

Here we develop a Bayesian statistical spatiotemporal model ensemble to interpret species-specific trends in the frequency of two target-site resistance mutations in the *Vgsc* gene, 995S and 995F, in three vector species *An. gambiae, An. coluzzii*, and *An. arabiensis* over the period 2005-2017, which encompasses the period of major scaling up of ITN distributions. The models are informed by 2418 observations of the frequency of each mutation in field sampled mosquitoes collected from 27 countries spanning western and eastern regions of Africa. For nine focal countries, we develop a series of fine resolution annual predictive maps. These models reveal the geographically structured patterns of spread of each mutation at both regional and continental scales. We use our geospatial predictions of *Vgsc* allele frequencies to address two questions of importance to malaria vector control. Firstly, we analyse associations between the *Vgsc* allele frequencies and phenotypic resistance to pyrethroids seen in field vector populations. Secondly, we explore the sensitivity of the predicted *Vgsc* allele frequencies to differences in the coverage of ITNs.

## RESULTS

### Predictive accuracy of the spatiotemporal model ensemble

Our spatiotemporal model ensemble, based on field-sampled *Vgsc* resistance allele frequencies in mosquito species from the African *An. gambiae* complex, confirmed our ability to interpolate allele frequencies. Predictive accuracy was assessed by testing the ability of the model ensemble to predict withheld data (using 10-fold out-of-sample cross-validation; see Methods), which showed a mean absolute prediction error (MAE; the average absolute difference between model predictions and observations) of less than 10% (MAE=0.083) across all observed *Vgsc* allele frequencies (with a root mean square error (RMSE) of 0.137; Supplementary Table S1).

### Spatiotemporal trends in the frequency of target-site resistance mutations

The nine mapped countries were chosen based on their number and spatial coverage of sampled *Vgsc* allele frequencies (see Methods and Supplementary Figures S1-S4). In western Africa, we developed maps of the predicted frequency of the *Vgsc*-995F mutation for Burkina Faso, Benin, Cameroon and Equatorial Guinea. In 2005, the earliest year in the data set, our maps show substantial geographic variation in the *Vgsc*-995F frequency, within each country and between countries. The marker frequency also varied markedly across the three vector species (Figure 1). In all four countries the marker frequency in 2005 was highest in *An. gambiae* and lowest in *An. arabiensis*, with frequencies in *An. coluzzii* also being low in large parts of each country. In Burkina Faso, Benin and Cameroon the marker frequency in 2005 is higher in southern compared to northern areas. We note that in these three countries the relative abundance of *An. arabiensis* declines southwards with decreasing latitude, with *An. gambiae* and *An. coluzzii* becoming more dominant (see Supplementary Figure S5). It is possible that there is a greater selection pressure for the development of insecticide resistance acting on *An. gambiae* and *An. coluzzii* populations, because these two species have a stronger tendency towards indoor human biting than *An. arabiensis* and are therefore more likely to encounter insecticide-treated surfaces (see the Discussion).

**Figure 1.**
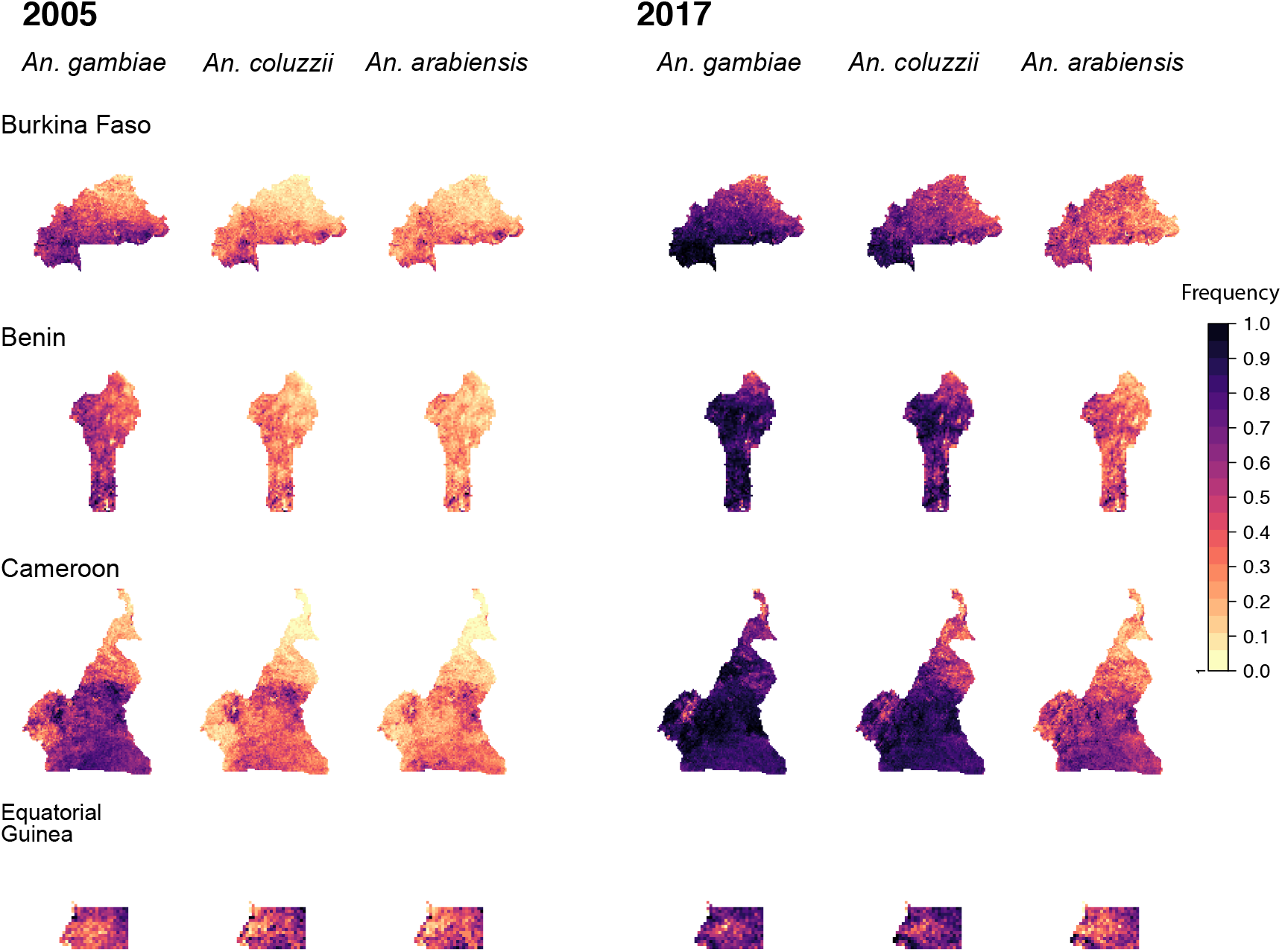
Predicted frequencies of the *Vgsc*-995F allele in malaria vector species for four countries in west Africa: Burkina Faso (top row), Benin (second row), Cameroon (third row), Equatorial Guinea (bottom row). Allele frequency maps for the first and final year are shown: the year 2005 is shown on the left (columns 1, 2 and 3) and the year 2017 is shown on the right (columns 4, 5 and 6). Columns 1 and 4 show maps for *An. gambiae*, columns 2 and 5 show maps for *An. coluzzii* and columns 3 and 6 show maps for *An. arabiensis*.

In all three vector species and all four countries, *Vgsc*-995F increased markedly between 2005-2017, with frequencies in *An. gambiae* and *An. coluzzii* in 2017 exceeding 0.5 in over 80% of the spatial area of each country (Figure 1). A lesser increase occurred in *An. arabiensis*, with the strongest rise occurring in southern Cameroon. We did not map the *Vgsc*-995S frequency for the countries in western Africa, owing to its general scarcity (full reasons for exclusion of countries from each part of the modelling analyses are provided in Table S2 in the Supplementary Material).

In eastern Africa, we developed maps of the predicted frequencies of *Vgsc*-995S and *Vgsc*-995F for four countries: Sudan, Ethiopia, Kenya and Uganda (Table S2). For Sudan, we mapped only a region in the west of the country (see Methods). The frequency of the *Vgsc*-995S allele in the four eastern African countries shows a dichotomous pattern across species, with much higher frequencies in *An. gambiae* than in *An. arabiensis* (Figure 2). In 2005, the frequency was low in *An. arabiensis* and very heterogeneous in *An. gambiae*. The frequency increased markedly in *An. gambiae* over 2005-2017, reaching very high levels in the north-west part of our mapped region in Sudan, south-east Ethiopia, west Kenya and most of Uganda. The *Vgsc*-995S frequency also increased in *An. arabiensis*, but to a much lesser extent, with the highest frequencies occurring in southern Uganda in the final year of the modelled time period.

**Figure 2.**
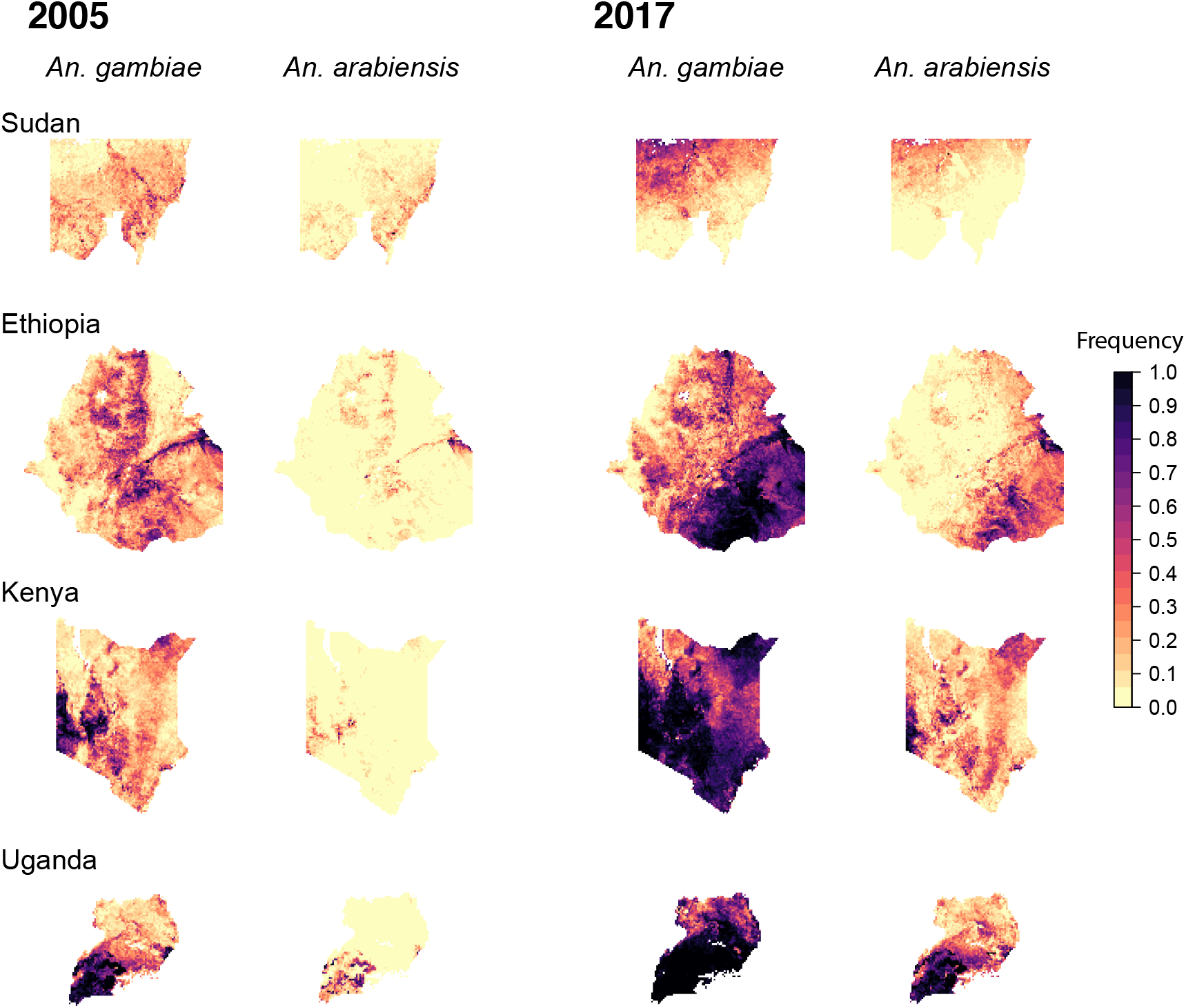
Predicted frequencies of the *Vgsc*-995S allele in malaria vector species for four countries in east Africa: Sudan (top row; the mapped area is confined to a region in the west (see Methods)), Ethiopia (second row), Kenya (third row), Uganda (bottom row). Allele frequency maps for the first and final year are shown: the year 2005 is shown on the left (columns 1 and 2) and the year 2017 is shown on the right (columns 3 and 4). Columns 1 and 3 show maps for *An. gambiae*, columns 2 and 4 show maps for *An. arabiensis*.

It is important to note that, in these four eastern African countries, the abundance of *An. arabiensis* relative to that of *An. gambiae* is typically much higher than in western Africa (Supplementary Figure S5), and *An. coluzzii* is very rarely reported. In Ethiopia and Sudan the species composition is almost entirely dominated by *An. arabiensis*. In general, the temporal dynamics of *Vgsc*-995F in *An. arabiensis* in these four countries followed a similar pattern to those in western Africa, with the frequency in 2005 being lower than that in *An. gambiae*, and then increasing in some areas to reach moderate to high frequencies in 2017 (Figure 3).

**Figure 3.**
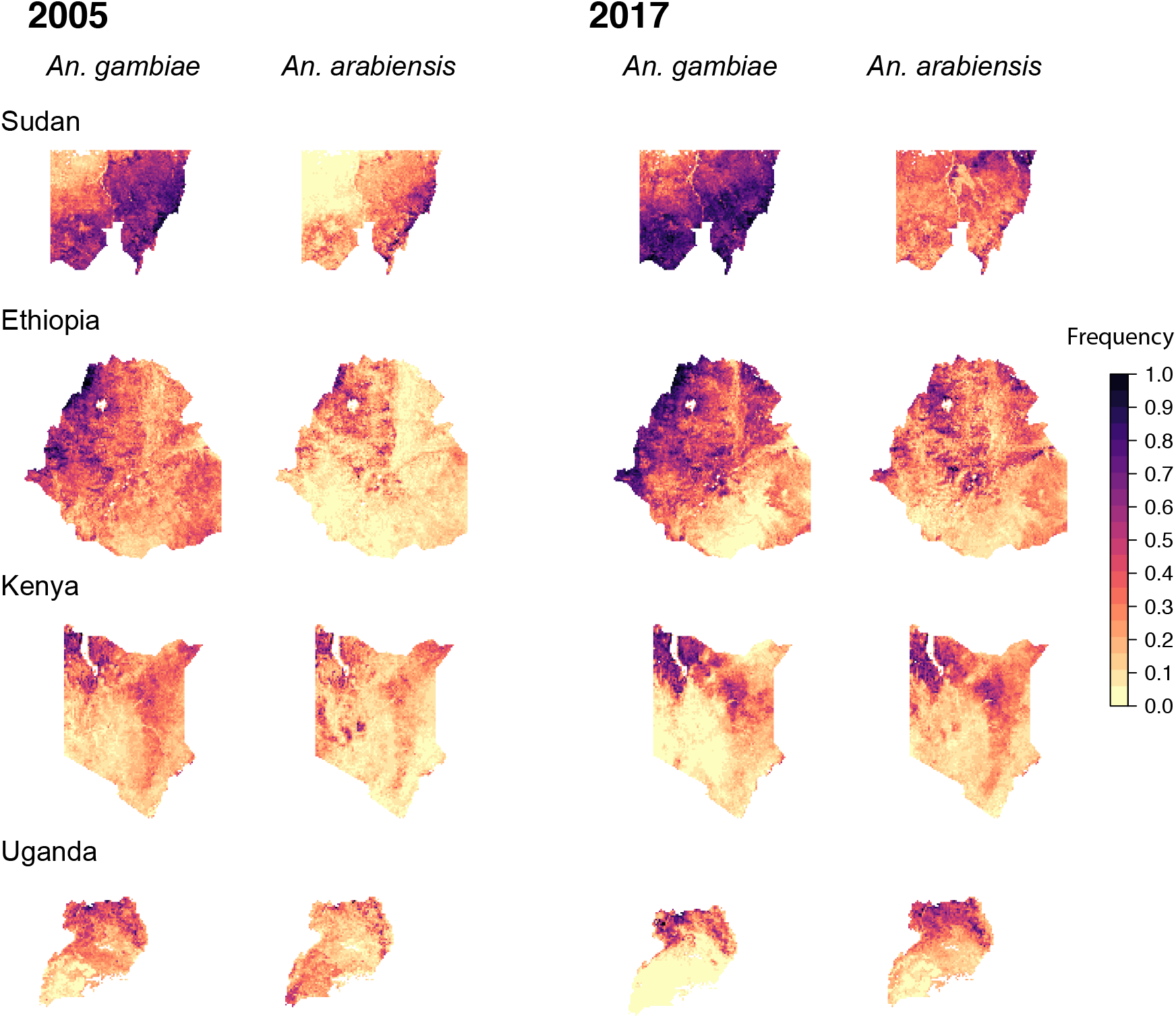
Predicted frequencies of the *Vgsc*-995F allele in malaria vector species for four countries in east Africa: Sudan (top row; the mapped area is confined to a region in the west (see Methods)), Ethiopia (second row), Kenya (third row), Uganda (bottom row). Allele frequency maps for the first and final year are shown: the year 2005 is shown on the left (columns 1 and 2) and the year 2017 is shown on the right (columns 3 and 4). Columns 1 and 3 show maps for *An. gambiae*, columns 2 and 4 show maps for *An. arabiensis*.

In Ethiopia, Kenya and Uganda, the frequency of *Vgsc*-995F in *An. gambiae* was typically lower in 2005 compared to the western countries, and there was a lesser increase in the frequency over 2005-2017 (Figure 3). In 2017 there was still substantial spatial heterogeneity in the *Vgsc*-995F frequency, with regions of high frequency in northwest Ethiopia, northwest Kenya and northern Uganda, and low frequencies elsewhere. In *An. gambiae*, the historical presence of *Vgsc*-995S at moderate to high frequencies (Figure 2) is likely to slow the spread of *Vgsc*-995F in this species (see the Discussion). In the south and west of our mapped region in Sudan, however, the *Vgsc*-995F frequency in *An. gambiae* was already high in 2005. Frequencies increased from 2005-2017, particularly in the north-western part of the region. For all four countries, there is a high degree of spatial overlap in the areas of relatively high *Vgsc*-995F frequency between *An. gambiae* and *An. arabiensis* (Figure 3).

For the DRC, we developed maps of the frequency of *Vgsc*-995F in *An. gambiae* only (Table S2). In the DRC, the spatiotemporal trends in *Vgsc*-995F in *An. gambiae* are more similar to the western countries, with a moderate to a high initial frequency in 2005, followed by a widespread increase to high frequencies in 2017 (Figure 4).

**Figure 4.**
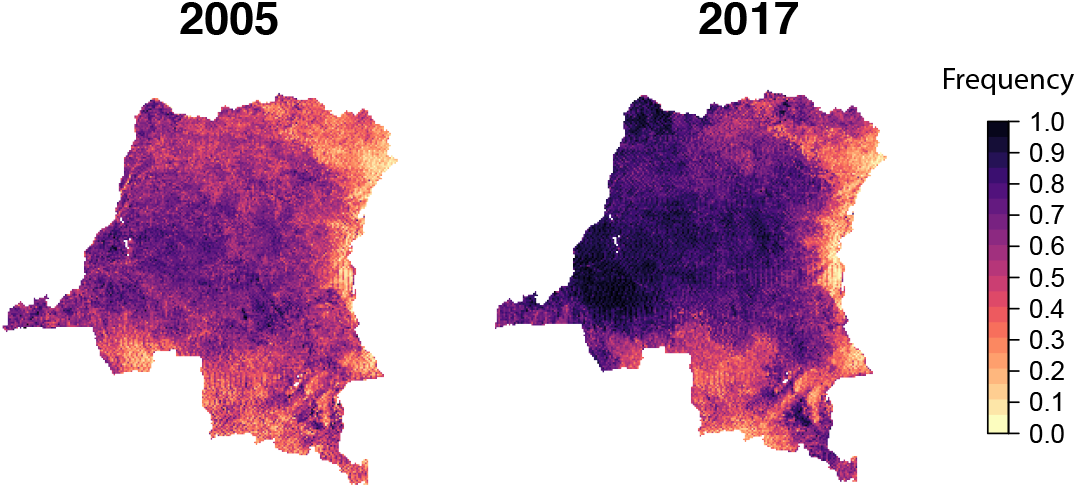
Predicted frequencies of the *Vgsc-*995F allele in *An. gambiae* in the Democratic Republic of the Congo (DRC). Allele frequency maps for the first and final year are shown: 2005 (left) and 2017 (right).

### Associations among the allele frequencies in the three vector species

The spatial patterns in the increases in *Vgsc*-995F frequencies in *An. gambiae* and *An. coluzzii* in the western countries over 2005-2017 were closely associated with each other, with the increase in *An. coluzzii* lagging behind that in *An. gambiae* (Figure 5). This is consistent with the results of genomic studies that show introgression of target-site resistance from *An. gambiae* to *An. coluzzii* (Clarkson *et al*. 2014). We found significant but less strong associations between the spatial patterns in *Vgsc*-995F frequency in *An. arabiensis* and both *An. gambiae* and *An. coluzzii* in the western countries over the years 2005-2017 (Supplementary Figure S6). Moreover, in the eastern countries (Ethiopia, Kenya, Uganda and Sudan) spatial increases in both the *Vgsc*-995F and *Vgsc*-995S frequencies were significantly associated across *An. gambiae* and *An. arabiensis* (Supplementary Figures S7 & S8).

**Figure 5.**
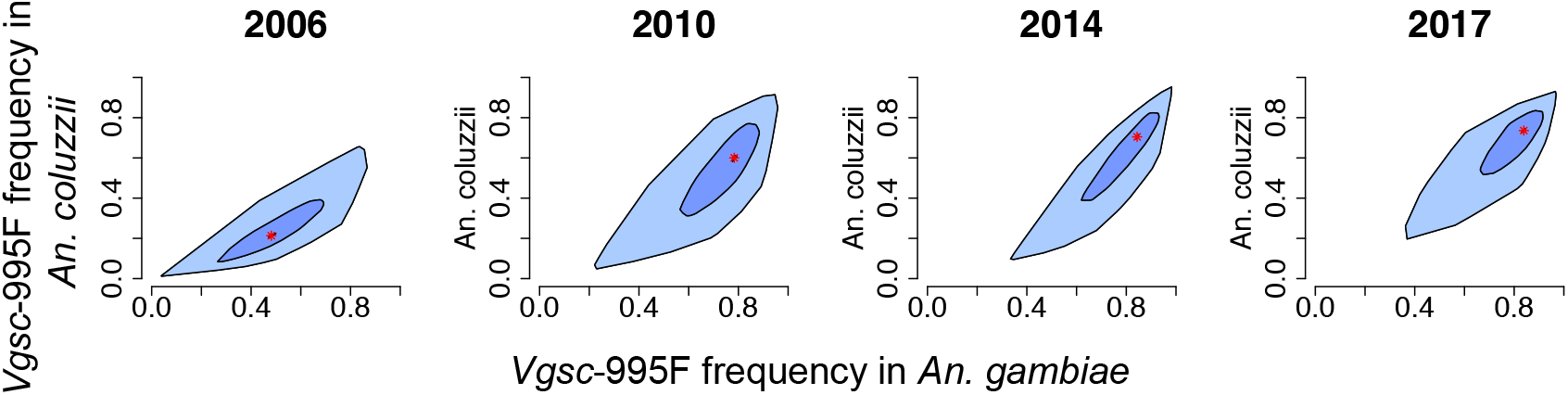
Associations between the predicted frequency of the *Vgsc*-995F allele in *An. gambiae* and *An. coluzzii*. Bagplots show the distribution across all mapped pixels within four countries in west Africa: Burkina Faso, Benin, Cameroon and Equatorial Guinea. The red asterisk shows the median, the dark blue shaded area contains 50% of all data points and the light blue shaded area contains all data points. Plots for four years are shown (from left to right): 2006, 2010, 2014 and 2017.

### Associations between resistance allele frequencies and the prevalence of resistance phenotypes

We investigated whether the variation in our mapped *Vgsc* mutation frequencies could explain variation in phenotypic resistance to pyrethroids in field malaria vector populations. Specifically, we analysed associations between predicted frequencies of the *Vsgc*-995F mutation in the mosquito samples and phenotypic resistance to deltamethrin, the most commonly used insecticide in malaria vector control during the period studied. Measures of mosquito mortality following exposure to deltamethrin were derived from standardised insecticide susceptibility tests (see Methods). We excluded Equatorial Guinea, Uganda, Kenya and the DRC from this analysis (Table S2). We do not consider associations between *Vsgc*-995S frequencies and the prevalence of deltamethrin resistance because *Vsgc*-995S frequencies are low in the majority of our selected countries and strongly segregated across the *An. gambiae* complex species (Figure 2 and see the Discussion).

For three countries in western Africa (Burkina Faso, Benin and Cameroon) and two countries in eastern Africa (Ethiopia and Sudan) the mortality to deltamethrin is consistently high when the *Vgsc*-995F frequency is close to zero, and there is a trend of decreasing mean mortality to deltamethrin with increasing *Vgsc*-995F frequency (Figures 6A & B). For each country, we fitted ordinary least-squares (OLS) linear regression models to the mean mortality values using the predicted *Vgsc*-995F frequency as a covariate (see Methods). The relationship with the *Vgsc*-995F covariate was significant for all countries except Sudan, in which case the 95% credible interval (CI) had a borderline overlap with zero (Table 1).

**Table 1.** OLS regression model results for each country. The model is fitted to mean mortality to deltamethrin across sets of bioassay sampling locations using the frequency of the *Vgsc*-995F allele in the Gambiae Complex as a covariate. The asterisk denotes statistical significance assessed by the 95% credible interval (CI).

| Data set | Intercept (95% CI) | *Vgsc*-995F (95% CI) | Adjusted $R^2$ | Degrees of freedom (df) |
| --- | --- | --- | --- | --- |
| Burkina Faso | 1.37*<br>(1.06, 1.67) | -0.65*<br>(-0.8, -0.51) | 0.34 | 157 |
| Benin | 2.1*<br>(1.1, 3.1) | -0.6*<br>(-0.1, -0.23) | 0.07 | 294 |
| Cameroon | 1.5*<br>(1.33, 1.66) | -0.54*<br>(-0.70, -0.38) | 0.33 | 182 |
| Ethiopia | -0.27 | -0.8* | 0.28 | 132 |
|  | (-0.59, 0.05) | (-1.0, -0.55) |  |  |
| Sudan | 1.1*<br>(0.85, 1.4) | -0.3<br>(-0.65, 0.04) | 0.03 | 254 |

**Figure 6.**
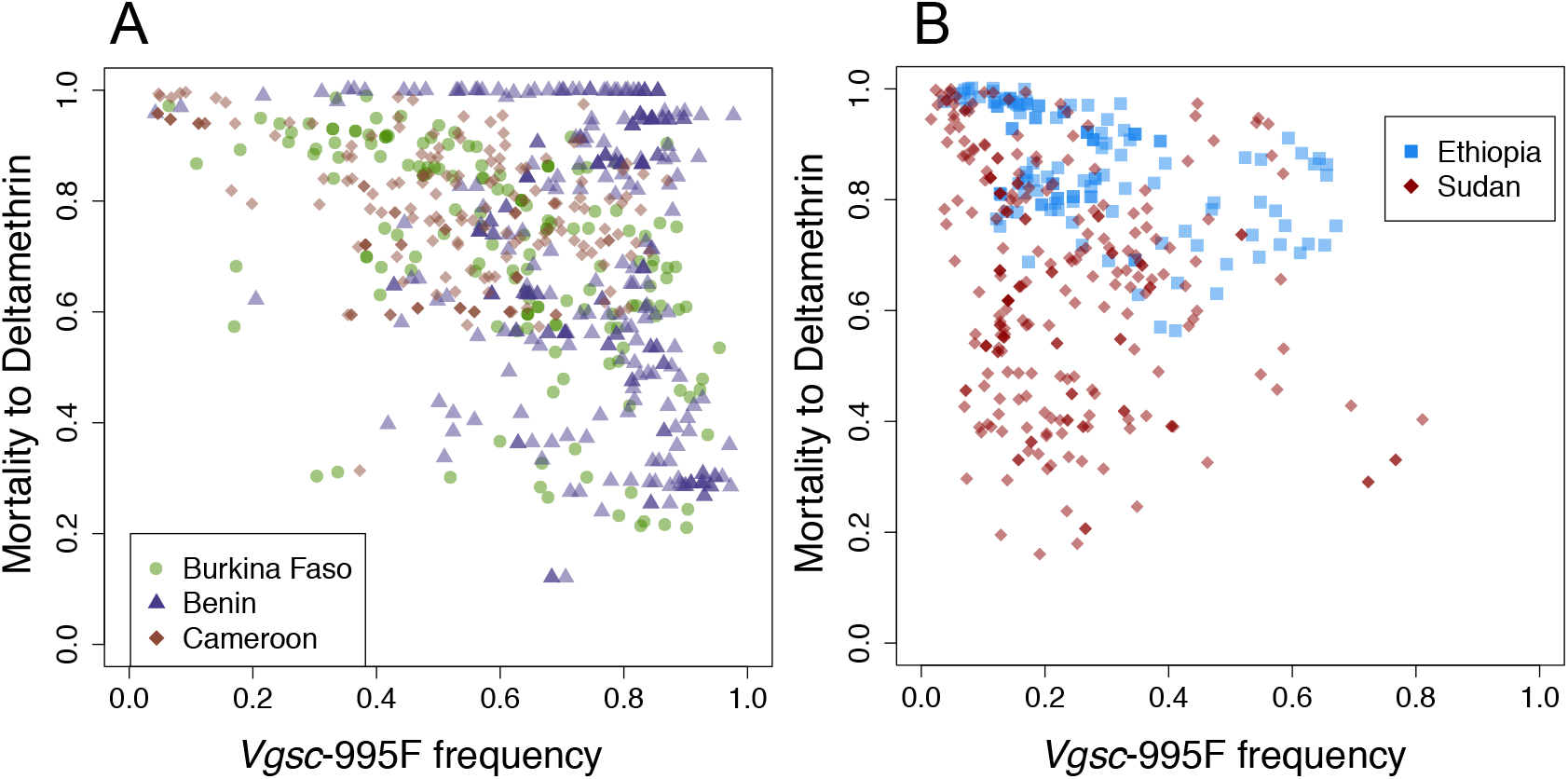
The mean mortality to deltamethrin and the frequency of the *Vgsc*-995F allele in Gambiae Complex mosquitoes at sampled locations in countries in west Africa (A) and east Africa (B). In the west (A), results for three countries are shown: Burkina Faso (green dots; *n*=159), Benin (purple triangles; *n*=297) and Cameroon (brown diamonds; *n*=184). In the east (B), results for two countries are shown: Ethiopia (blue squares; *n*=134) and Sudan (dark red diamonds; *n*=256).

Despite the uncertainty associated with estimating frequencies of both phenotypic resistance and *Vgsc* alleles across multiple mosquito species in field populations, the interpolated *Vgsc*-995F allele frequency is able to partially explain the variation in mortality to deltamethrin. The level of explained variation varies across countries; adjusted *R*^2^ values were close to 0.3 for Burkina Faso, Cameroon and Ethiopia, but less than 0.1 for Benin and Sudan (Table 1). Moreover, the form of the relationship varies across countries. In Benin many mortality values remain high across increasing *Vgsc*-995F frequencies (Figure 6), consistent with the poor explanatory value of the model, despite a significant negative slope (Table 1).

### Relationships with predictor variables

Our model ensemble included 99 predictor variables describing environmental and biological processes that could potentially drive selection for insecticide resistance (see Methods). We analysed which of these variables were the most influential predictors of *Vgsc* allele frequencies using variable importance measures, which describe the influence of each variable in terms of its impact on model predictions, relative to all other predictor variables (see Methods). Our model ensemble included three constituent models: an extreme gradient boosting model (XGB), a random forest model (RF) and a neural network model (NN). For each model we obtained a ranking of the most influential variables using a variable importance measure that was chosen based on the type of model (see Methods).

For all three models, the highest-ranked predictor variable was related to climate, with solar radiation ranking highest for the XGB and RF models and relative humidity ranking highest for the NN model (Table 2). These two variables may be influential because they segregate dry arid areas and wetter tropical regions (see the Discussion). The coverage of Insecticide Treated Bednets (ITNs) was strongly influential in the XGB and RF models, with variables describing ITN coverage at different time lags ranking second, fifth and ninth in both models (Table 2). In the NN model, the coverage of evergreen broadleaf forest was highly influential, with different time lags of this variable ranking second, fourth and eighth. In general, with the exception of ITN coverage, the highest-ranked variables for the XGB and RF models are related to climate and elevation, and the highest ranked variables for the NN model include variables relating to land cover, climate and elevation.

**Table 2.** The top ten highest ranked variables, as determined by variable importance measures, for the three machine learning models included in the model ensemble. Variable name suffixes (−1), (−2), and (−3) denote time lags of 1, 2, and 3 years, respectively. One, two, and three asterisks denote the first, second, and third principal component, respectively, for variables available on a monthly time step (see Methods).

| Rank | XGB | RF | NN |
| --- | --- | --- | --- |
| 1 | Solar radiation*** | Solar radiation*** | Relative humidity* |
| 2 | ITN coverage (-1) | ITN coverage (-1) | Evergreen broadleaf forest (-3) |
| 3 | Elevation | Elevation | Wind speed* |
| 4 | Daytime temperature**(-2) | Tassel cap brightness**(-2) | Evergreen broadleaf forest (-1) |
| 5 | ITN coverage (-2) | ITN coverage (-2) | Elevation |
| 6 | Night time temperature* (-2) | Wind speed* | Cropping factor |
| 7 | Enhanced vegetation index**(-1) | Tassel cap brightness**(-3) | Cation exchange capacity |
| 8 | Rainfall*(-3) | Temperature diurnal difference*(-2) | Evergreen broadleaf forest (-2) |
| 9 | ITN coverage | ITN coverage | Tropical fruit |
| 10 | Night time temperature** (-2) | Tassel cap brightness** | Daytime temperature*(-1) |

### Rank XGB RF NN

#### Impacts of increasing ITN coverage on predicted *Vgsc* mutation frequencies

Relationships between ITN coverage and the development of insecticide resistance have significant implications for malaria vector control (see the Discussion). We further examined the relationship between ITN coverage and the interpolated frequencies of the *Vgsc*-995F allele by calculating the independent conditional expectation (ICE) of the predicted frequency under changing ITN coverage (Goldstein *et al*. 2015; Lucas 2020). The ICE can be calculated for any of the locations (pixels) of our predictive maps, and we selected a single location in each country to evaluate the ICE (see Methods). We chose to analyse these relationships for the year 2005, because up until this time the resistance allele frequencies were unlikely to be affected by widespread ITN usage (the reported ITN coverage in this year and the three years prior is very close to zero). We varied the ITN coverage in the years 2002-2005 from zero to 0.9, or 0-90% of people slept under an ITN the preceding night, in increments of 0.1 (because the predictor variables include three annual time lags; see Methods). It is important to note that this variation in ITN coverage that we simulate was never actually observed in the period 2002-2005. We did not analyse relationships between ITN coverage and the *Vgsc*-995S allele frequency because *Vgsc*-995S shows low frequencies for all years in most of the countries included in our model.

For the four countries in western Africa, increasing the ITN coverage causes the model to predict increasing *Vgsc*-995F frequencies in 2005 in all three mosquito species (Figure 7A). The impact of increasing ITN coverage varies geographically across countries, and also between mosquito species. In *An. gambiae*, the *Vgsc*-995F frequency at the selected location was already high (>0.4) in 2005 for all countries except Benin. Increasing the ITN coverage from zero to 0.9 resulted in further increases in frequencies to very high values, with the largest increases occurring when the frequency at zero ITN coverage was lower (Figure 7A). In *An. coluzzii, Vgsc*-995F frequencies in 2005 were relatively low at zero ITN coverage, and predicted frequencies increased strongly as ITN coverage increased to high values. *An. arabiensis* showed the lowest *Vgsc*-995F frequencies in 2005 under zero ITN coverage, and the impact of increasing ITN coverage on predicted frequencies was much less than in *An. gambiae* and *An. coluzzii*. This behaviour is consistent with the trends in *Vgsc*-995F in *An. arabiensis* over the years 2005-2017 (Figure 1), with frequencies remaining relatively low while the coverage of ITNs increased from 2005 onwards in all four countries to reach moderate to high values (Supplementary Figures S9-S11).

**Figure 7.**
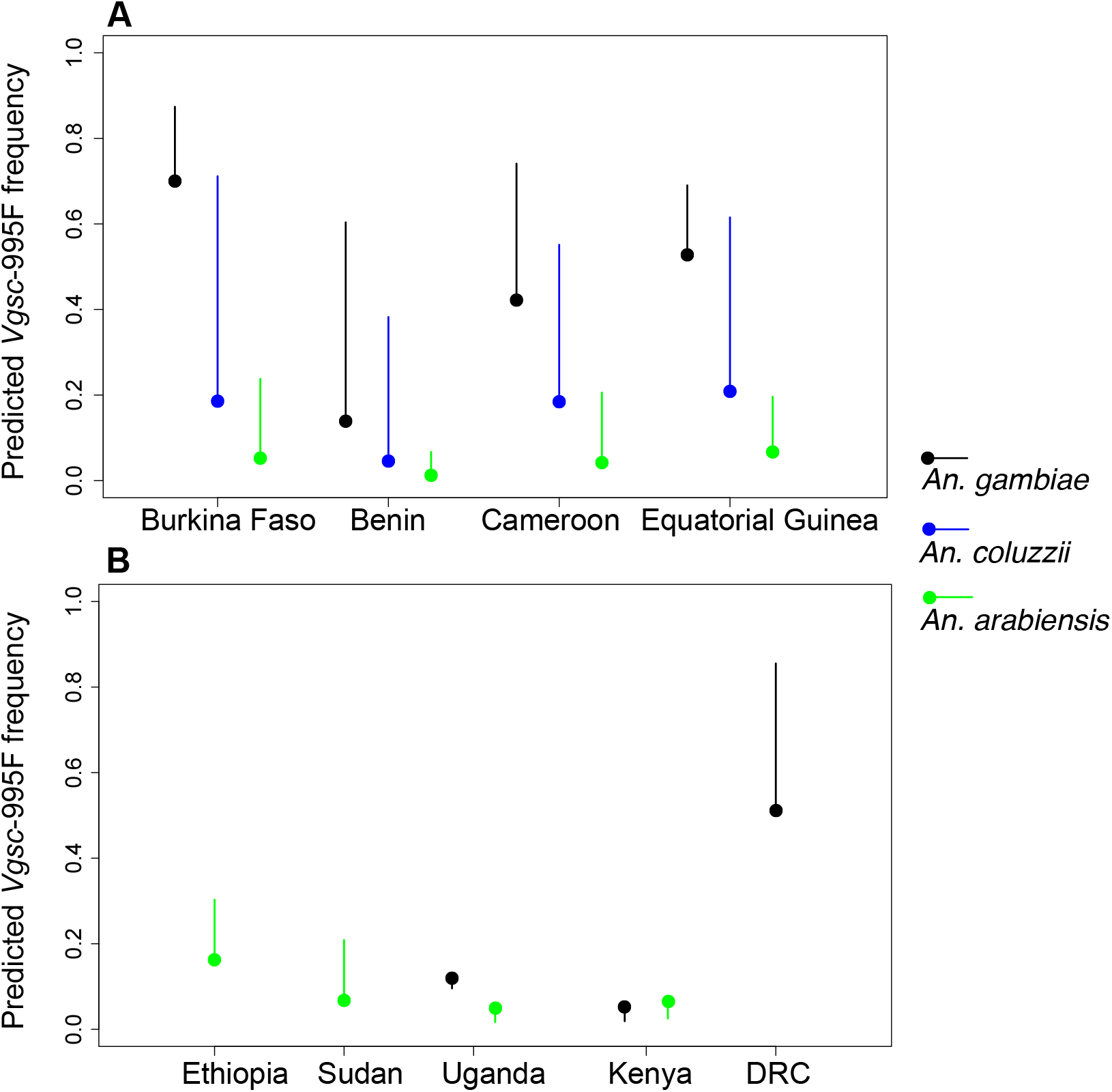
The variation in the model-predicted *Vgsc*-995F frequency in malaria vector species for the year 2005 as the coverage of insecticide treated bednets (ITNs) is increased. Within each country, the predicted frequency for a single point location is shown (see text). Solid circles represent the predicted frequency corresponding to an ITN coverage of zero over the years 2002-2005, which is close to the recorded ITN coverage over this period. Solid lines show the range of variation in the predicted frequency at these locations for the year 2005 as the ITN coverage is increased from 0-0.9. Results for four countries in west Africa (**A**) and five countries in central and east Africa (**B**) are shown. Black, blue and green lines and circles represent predicted frequencies in *An. gambiae, An. coluzzii*, and *An. arabiensis*, respectively.

For the four countries in eastern Africa, the impact of increasing ITN coverage on the model predictions of *Vgsc*-995F frequencies is relatively small (Figure 7B), reflecting the differences in malaria vector species composition, and the different types of insecticide resistance mechanisms present, between the eastern and western regions of Africa (see the Discussion). In Ethiopia and Sudan, the species composition consists mostly of *An. arabiensis* (see Supplementary Figure S5); the *Vgsc*-995F frequencies in *An. arabiensis* were low in 2005 and increasing ITN coverage resulted in only a small increase in frequencies. In Kenya and Uganda, the predicted *Vgsc*-995F frequencies were almost unchanged by increasing ITN coverage. This is consistent with the trends in *Vgsc*-995F frequencies in both *An. arabiensis* and *An. gambiae* in eastern Africa from 2005-2017 (Figure 3), with *Vgsc*-995F frequencies increasing by a small amount in some areas and not increasing at all in other areas, although ITN coverage did increase in these countries over the period (Supplementary Figures S9-S11). In *An. gambiae*, the earlier increases in the frequencies of *Vgsc*-995S may have reduced the selection pressures driving the spread of the *Vgsc*-995F allele (see the Discussion). In the DRC, the variation in predicted *Vgsc*-995F frequencies with increasing ITN coverage shows a similar pattern to the western countries, with predicted frequencies in *An. gambiae* in 2005 increasing from an intermediate value at zero ITN coverage to a very high value with increasing ITN coverage (Figure 7B).

## DISCUSSION

Our annual maps of the frequencies of target-site insecticide resistance mutations in the three dominant malaria vector species of the African *An. gambiae* complex have characterised the species-specific spread dynamics of target-site resistance, showing how these dynamics have varied geographically, across national and continental scales. Our geospatial machine learning model ensemble brings together data sets describing multiple, multifaceted processes affecting insecticide resistance in field vector populations. Relationships between mutation frequencies and the prevalence of resistance phenotypes can be explored, as well as relationships with the coverage of vector control interventions such as ITNs. In this study, we have used these models to investigate questions of importance to malaria vector control.

Firstly, we found significant relationships between frequencies of the target-site resistance mutation *Vgsc-*995F and phenotypic resistance to the pyrethroid deltamethrin in field samples. This demonstrates explanatory power of target-site resistance for phenotypic variation in field *An. gambiae* complex populations, supporting the relationships between target-site and phenotypic resistance shown by functional (Ranson *et al*. 2000; Grigoraki *et al*. 2021) and genomic (Miles *et al*. 2017b; Clarkson *et al*.) studies. Our maps show substantial spatial heterogeneity in *Vgsc* allele frequencies in recent years, with frequencies in 2017 varying both across vector species and geographic regions. Continued surveillance of these target-site markers is therefore important to track current and future regional temporal trends in resistance.

A substantial amount of phenotypic variation was unexplained by the *Vgsc*-995F frequencies, however, which is in part due to the fact that the sample mortality is often not disaggregated by the individual species within the *An. gambiae* complex. Our maps show how target-site resistance frequencies differ across these species depending on geographic region, with dichotomous differences in some regions. This highlights that species-specific trends in phenotypic resistance cannot be fully understood using susceptibility test mortality values at the *An. gambiae* complex level. According to our *Vgsc* allele frequency maps, in Kenya and Uganda there is a close coupling between vector species and which type of target-site resistance mechanism is more prevalent (995F or 995S), therefore errors due to aggregating across species will be particularly high in these countries. For this reason, we were unable to assess associations between phenotypic pyrethroid resistance and frequencies of the *Vgsc-*995S mutation, which has a high frequency in these two countries and relatively low frequencies in our other focal countries.

In addition to target-site resistance, geospatial analysis of the distribution of metabolic resistance mechanisms could greatly improve our ability to understand spatiotemporal trends in resistance. Metabolic resistance, the insect’s increased ability to metabolise insecticide, is another important mechanism that can generate high levels of pyrethroid resistance in *An. gambiae* (Edi *et al*. 2014; Mitchell *et al*. 2014), and especially *Anopheles funestus*, which lacks resistance-associated *Vgsc* mutations (Riveron *et al*. 2019). Metabolic resistance occurs through the upregulation of metabolic genes that encode detoxification enzymes. Many metabolic genes have shown associations with pyrethroid resistance, with genomic studies of the *An. gambiae* complex finding strong signals of positive selection around gene clusters implicated in insecticide metabolism (Miles *et al*. 2017b; Njoroge *et al*. 2021). Amplicon sequencing panels, which screen a panel of markers of interest across many loci (Lucas *et al*. 2019; Makunin *et al*. 2021), can incorporate target-site as well as metabolic resistance markers, including known mutations in the *Gtse2* and *Cyp6p* gene clusters (Miles *et al*. 2017b). The anticipated increased use of amplicon sequencing panels in genetic surveillance of vector populations in the coming years will lead to exciting opportunities to better quantify the polygenic nature of resistance.

Secondly, our models indicate that the coverage of ITNs is influential in predicting *Vgsc* allele frequencies, but that the strength of this influence varies both geographically and across vector species. These relationships between ITN coverage and *Vgsc* allele frequencies produced by our models need to be interpreted with caution, because the machine learning approaches that we have applied do not allow causal inferences to be made, and correlations amongst predictor variables can make relationships with any given variable difficult to identify. Nonetheless, our results are consistent with evidence from field studies showing changes in *Vgsc* allele frequencies following the implementation of ITN interventions. Our results showed that ITN coverage had the greatest influence on *Vgsc-*995F frequencies in the western African countries in *An. gambiae* and *An. coluzzii*. Field studies have also shown increases in *Vgsc-*995F frequencies in these two species following the scale-up of ITNs in Cameroon (Mandeng *et al*. 2019), Ghana (Lynd *et al*. 2010) and Mali (Norris *et al*. 2015). In Kenya and Uganda, we found no influence of ITN coverage on predicted *Vgsc-*995F frequencies in *An. gambiae*, which reflects the more limited spread of the 995F mutation in eastern *An. gambiae* populations. It is possible that the spread of *Vgsc-*995F in *An. gambiae* in eastern Africa was inhibited by the presence of the *Vgsc-*995S mutation, which is known to have been present in Kenyan *An. gambiae* populations since 1986 (Ranson *et al*. 2000). Resistance conferred by *Vgsc*-995S could lead to reduced selection for *Vgsc-*995F, and it is also possible that the strength of selection could have been reduced if other mechanisms, such as metabolic resistance, were already present.

The influence of ITN coverage on predicted allele frequencies is consistently lower in *An. arabiensis* across all nine countries compared to the other two species, which reflects the more limited spread of both *Vgsc* alleles in *An. arabiensis* across the western and eastern countries. *An. arabiensis* have a greater tendency towards biting outdoors than *An. gambiae* and *An. coluzzii*, and their peak biting times occur earlier in the evening while the other two species bite most commonly in the middle of the night (Fornadel *et al*. 2010; Sinka *et al*. 2010; Russell *et al*. 2011). *An. arabiensis* also has a lower human blood index than *An. gambiae* and *An. coluzzii*, indicating a relatively high proportion of bites taken on animals rather than humans (Mayagaya *et al*. 2015). It is therefore plausible that *An. arabiensis* has lower exposure to ITNs, and thus ITN coverage has a lesser impact on selection for resistance in this species. Observed shifts in vector species composition towards higher proportions of *An. arabiensis* following the scaling up of ITN interventions supports this hypothesis (Russell *et al*. 2011; Sinka *et al*. 2016). The evolutionary pathway of resistance differs across the three vector species, however, and we expect greater divergence in the case of *An. arabiensis* which has lower rates of hybridisation with the other two species (Weetman *et al*. 2012), with higher rates of hybridisation occurring between *An. gambiae* and *An. coluzzii* (Vicente *et al*. 2017). For example, hybridisation led to the introgression of target-site resistance from *An. gambiae* to *An. coluzzii* (Clarkson *et al*. 2014; Norris *et al*. 2015), accelerating the development and spread of target-site resistance in *An. coluzzii*.

Variables describing solar radiation and humidity were the highest ranking in terms of their impact on predicted allele frequencies. While we cannot identify a mechanistic explanation for this result, we note that these climate variables provide a broadscale spatial separation of areas that are arid from those that are wet and tropical. They may, therefore, represent unmeasured differences in mosquito population structure and genetics that give rise to regional differences in resistance patterns. We also found, however, that the most influential variables were different across the different machine learning models, with the neural network model showing less commonality with the two regression tree-based models (extreme gradient boosting and random forest). Lucas et al. (2020) found that variable importance measures for neural network models are not replicable, with different variable importance rankings being produced each time the model is fitted to the same data set. This emphasises that models fitted by machine learning algorithms do not represent a single unique optimal solution, and there may be many different ways that a machine learning model can combine the predictor variables to produce similarly accurate results, as measured by out-of-sample testing (Hastie *et al*. 2009).

In conclusion, our geospatial analyses illustrate how insecticide target-site resistance dynamics in African malaria vectors vary across species and geographic regions, emphasising that resistance management strategies need to be based on local information about resistance genetics and vector species composition, as well as phenotype surveillance. Our results demonstrate that genetic surveillance of resistance can help to predict resistance phenotypes in field vector populations and understand their mechanistic drivers. This capacity would be improved by surveillance of resistance phenotypes at the level of individual vector species. In addition to target-site resistance, surveillance of other genetic resistance mechanisms, such as metabolic resistance, is needed to understand, predict and manage the spread of resistance.

## MATERIAL AND METHODS

### Summary

We analysed trends in the frequencies of target-site insecticide resistance mutations across space and time in three African malaria vector species: *An. gambiae, An. coluzzii* and *An. arabiensis*. We use spatiotemporal modelling approaches that apply both Bayesian statistics and machine learning methods in order to predict mutation frequencies jointly across the three species over a spatial grid of approximately 5 km resolution. Our model predictions are based on surveillance data that records observed frequencies in mosquitoes sampled widely throughout west and northeast Africa. The machine learning methods predict the proportions of each allele in each mosquito sample, and they are informed by 99 potential predictor variables that represent environmental and biological processes which may influence selection for resistance. A Bayesian multinomial metamodel then combines predictions across the multiple machine learning models in order to make more accurate and robust predictions (a methodology known as stacked generalisation (Wolpert 1992)). Using the metamodel we predict the frequencies of each mutation in all grid cells within our nine selected countries for all years in the period 2005-2017.

### *Vgsc* allele frequency data

Our models are informed by a database containing frequencies of *Vgsc* mutations in mosquito samples belonging to the *Anopheles gambiae* species complex collected from within western and eastern Africa over the period 2005–2017. This database is an updated version of a publicly available data set containing *Vgsc* allele frequencies (Moyes *et al*. 2019b) and collates data sets from multiple contributors, including published and unpublished sources. The database records the number of mosquitoes tested in each sample, together with the frequencies of the *Vgsc*-995L, *Vgsc*-995F and *Vgsc*-995S alleles in the sample. Some, but not all, data sets in the database record the *Vgsc* genotype of the sampled mosquitoes. The database also records information about the mosquito species tested, the molecular screening methods used for species identification and *Vgsc* allele identification, and the geographic coordinates of the sample collection location. We only included samples that are representative of the *An. gambiae* population sampled at each place and time (i.e. randomly sampled from the population). We also only included samples that contained five or more mosquitoes. The final data set included 2418 samples distributed across 27 countries.

We developed predictive maps of *Vgsc* allele frequencies for a focal selection of countries which had the highest number of samples, excluding those countries for which the spatial distribution of samples was strongly clustered (Figures S1 and S2). In selecting countries for inclusion in our mapping analysis, we subdivided the African continent into western and eastern regions, with Cameroon and countries further west of Cameroon falling within our western region and countries that lie east of the Central African Republic falling within our eastern region. Within the western region, we selected the five countries with the greatest number of samples (Supplementary Figure S3), excluding Senegal because of a tight clustering of sampling locations around the border with The Gambia (Figure S1). In the eastern region, we selected all countries that had samples that were included in our modelling analysis (Figure S4), excluding Tanzania due to a strong spatial clustering of the sampling observations (Figure S2). Sudan is the most data-rich country included in our study (Figures S3 and S4) but it covers a large spatial area and the sampling locations are all located in a region in the eastern part (Figure S2). Therefore, we developed predictive maps only for a region in the east of Sudan that does not extend further west than a longitude of 29.5°E or further north of 17°N. In the case of Ethiopia, we excluded the region east of a longitude of 44°E because we have no samples located in this region.

We included one central African country, the Democratic Republic of Congo (DRC), in our mapping analysis. Although the *Vgsc* allele frequency data is sparse throughout the country (Figures S1, S2 and S4), we included the DRC because it covers a region that is rarely studied. In the case of the DRC, our modelling analysis is restricted to predicting the frequency of the *Vsgc*-995F mutation only, and we do not predict *Vsgc*-995S frequencies (see below). We excluded the data on *Vsgc*-995S frequencies from the DRC analysis because most studies from the DRC only perform an assay capable of detecting L995F, which can lead to erroneous genotypes when both resistant alleles are present, which appears typical in DRC (Loonen 2020, Lynd et al. 2018).

### Potential predictor variables

Our set of predictors is similar to that described in Hancock et al. (Hancock *et al*. 2020; Moyes *et al*. 2020), and includes 99 variables describing environmental characteristics that could potentially be related to the development and spread of insecticide resistance in populations of *Anopheles gambiae* complex mosquito species. These variables describe the coverage of insecticide-based vector control interventions, agricultural land use (You *et al*. ; Friedl & Sulla-Menashe 2015), and the environmental fate of agricultural insecticides (Hendriks *et al*. 2019), other types of land use (Friedl & Sulla-Menashe 2015; Esch *et al*. 2017; Tatem 2017; Sulla-Menashe *et al*. 2019), climate (Trabucco & Zomer 2009; Friedl & Sulla-Menashe 2015; Funk *et al*. 2015), and relative species abundance. A detailed description of this set of predictor variables is provided in Table S3 of the Supplementary Material. Our vector control intervention data includes a variable estimating ITN coverage in terms of the proportion of people who slept under a net the preceding night, at each ∼5 km pixel location for each year (Bhatt *et al*. 2015b; Weiss *et al*. 2019). Relative species abundance is represented by a variable estimating the abundance of *An. arabiensis* relative to the abundance of *An. gambiae* and *An. coluzzii* (Sinka *et al*. 2016). For all variables, we obtained spatially explicit data on a grid with a 2.5 arc-minute resolution (which is approximately 5 km at the equator) covering sub-Saharan Africa. For variables for which temporal data were available at an annual resolution, we included time-lagged representations with lags of 0, 1, 2, and 3 years.

### Stacked generalization ensemble modelling approach

We used stacked generalization to develop a model ensemble that combines the predictions generated by multiple machine learning models (Wolpert 1992; Ting & Witten 1997). Stacked generalization uses a meta-model, or “generalizer”, that learns a weighted combination of the predictions across each model in the ensemble, where the predictions of each model are the out-of-sample predictions derived from *K-*fold cross validation. The predictions produced by the generalizer correct for the biases of each model, and are expected to have improved prediction accuracy relative to any of the individual models included in the ensemble (Wolpert 1992; Ting & Witten 1997; Bhatt *et al*. 2017).

#### Machine learning models

Our model ensemble included three different machine learning models that predicted the frequencies of the *Vgsc*-995L, *Vgsc*-995F and *Vgsc*-995S at each pixel within our mapped countries for each year within the period 2005-2017. The three machine learning models were an extreme gradient boosting (XGB) model, a random forest (RF) model and a neural network (NN) model. These models were chosen due to their demonstrated high predictive performance (Crisci *et al*. 2012; Bhatt *et al*. 2017), which derives from their ability to represent non-linear relationships and high-level interactions across the model features (Bhatt *et al*. 2017; Hancock *et al*. 2020). The XGB model was implemented using the R package xgboost (Chen & Guestrin 2016) and the RF and NN models were implemented using the sklearn (Pedregosa *et al*. 2011) and keras packages (Chollet & others 2015) in Python. The label for these models was a categorical variable corresponding to whether the *Vgsc*-995L, *Vgsc*-995F or *Vsgc*-995S mutation was detected across all the alleles screened in each sample. All *Vgsc* allele frequency observations from the 27 countries in our data set were used to inform the model (see above). The models predict the expected frequencies of each allele at each mapped pixel. The features used in the models included the 99 environmental predictor variables together with the 1-, 2-, and 3-year lags for those variables that vary on a yearly time step. A factor variable representing the mosquito species (*An. gambiae, An. coluzzii, An. arabiensis*, or *An. gambiae s*.*l*) was also included as a feature, where the *An. gambiae s*.*l*. category describes individuals within samples for which species within the *Anopheles gambiae* complex were not identified. Finally, the year in which the bioassay and allele frequency samples were collected was also included as a feature. For each machine learning model, parameter tuning was performed using out-of-sample validation by subdividing the data into training, validation and test subsets (see the Supplementary Material).

We developed an additional model ensemble that predicted only the frequency of *Vsgc*-995F, which we used to develop predictive maps of the *Vsgc*-995F frequency for the DRC. This model ensemble included the three machine learning models as described above, and the label was a categorical variable corresponding to whether the *Vsgc*-995F mutation was detected across all the alleles screened in the sample. The label included the full data set containing the *Vsgc*-995F frequencies in the 2418 samples. The features used were the same as those used in the above models, and parameter tuning was performed as described above.

#### Model stacking and multinomial logistic regression

We use a Bayesian multinomial logit regression model as our meta-model to combine the out-of-sample predictions obtained from performing *K-*fold cross-validation on each of the three machine learning models in the model ensemble (Baker 1994; Rue *et al*. 2009; Croissant 2010) (see www.r-inla.org). The multinomial logit model represents observations where the sampling unit corresponds to one of a set of mutually exclusive alternatives *j* ∈ {1, …, *J*}; in our case *J*=3, with the alternatives being the *Vgsc*-995L, *Vgsc*-995F or *Vgsc*-995S marker (we do not account for diploid genotypes in our model). Our observations *y*_*ij*_ are the numbers of *Vgsc*-995L alleles (*j*=1), *Vsgc*-995F alleles (*j*=2) and *Vsgc*-995S alleles (*j*=3) in sample *i*, with *i*=1,….,*N* samples in total. Our model has three covariates which are the out-of-sample predictions of the frequencies of each allele in each sample given by the three machine learning models, transformed using the empirical logit transform to avoid discontinuities at 0 and 1. We store these covariates in the matrices **X**^1^, **X**^2^, and **X**^3^, which have dimension *N*×*J*, with each matrix containing the predictions of frequencies of the three alleles for one of the three machine learning models. Our multinomial logit model uses the following linear predictor:

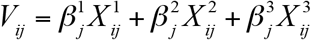

where there are three sets of three coefficients 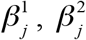 and 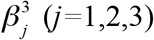; we combine these into the vector B. For each observation *i* the expected probabilities of each alternative are:

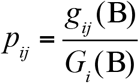

Where *g*_*ij*_ (B) = exp(*V*_*ij*_) and 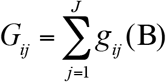 (see Croissant (2010) and www.r-inla.org).

We use the multinomial-Poisson transformation (Baker 1994), which gives the following expression for the Poisson likelihood (Baker 1994):

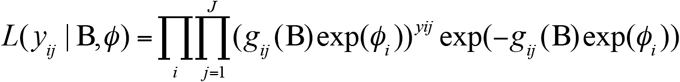

where *ϕ*_*i*_ are *N* additional parameters that need to be estimated in order to use the multinomial-Poisson transformation. Posterior distributions of the parameters B and *ϕ*_*i*_ are obtained by fitting the model using the R-INLA package (Rue *et al*. 2009) (see www.r-inla.org), with the coefficients B as fixed effects and the intercepts *ϕ*_*i*_ as an independent (iid) random effect. Our implementation constrains each of the nine coefficients to be positive 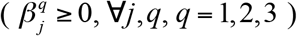 (Bhatt *et al*. 2017). Once the parameter estimation has been performed, the final set of predictions given by the model ensemble are obtained by replacing the elements of **X**^1^, **X**^2^, and **X**^3^ with the in-sample predictions of the machine learning models obtained by fitting each of these models to all the data (all the labels and the corresponding sets of features). For our second model ensemble for predicting only *Vsgc-*995F frequencies, the formulation of the meta-model is the same as described above, with *J*=2.

### Posterior validation

To assess the ability of our model to accurately represent the data, we performed posterior validation of our model ensemble using 10-fold out-of-sample cross-validation. Specifically, the data were divided into 10 subsets (or “test” sets, using random sampling without replacement), and 10 successive model fits were performed, each withholding a different test set. The test sets were withheld from each of the three machine learning models included in the ensemble, as well as from the multinomial logit metamodel. The root mean squared error (RMSE) across all (withheld) *Vgsc* allele frequency observations confirmed that the model ensemble delivered higher prediction accuracy than each of the three machine learning model constituents (Supplementary Table S1).

### Insecticide resistance bioassay data

To analyse relationships between our predicted resistance allele frequencies and resistance phenotypes observed in field vector populations, we utilised a database of insecticide resistance bioassay data (Hancock *et al*. 2020) including samples tested over the period 2005-2017. All species included in the samples are from the *Anopheles gambiae* complex and the composition of sibling species is unknown for the majority of samples. The data record the number of mosquitoes in the sample and the proportional sample mortality resulting from the bioassay, as well as variables describing the mosquitoes tested, the sample collection site, and the bioassay conditions and protocol. We selected the bioassay results for standard diagnostic dose WHO susceptibility tests performed using deltamethrin for all samples collected within the five countries included in our analysis (see Results), resulting in 159 results for Burkina Faso, 297 results for Benin, 184 results for Cameroon, 134 results for Ethiopia and 256 results for Sudan. The bioassay data set included only two bioassay results for Equatorial Guinea and 22 bioassay results for Uganda, so we excluded these countries from our analysis of associations between our mapped *Vgsc* allele frequencies and the prevalence of insecticide resistance phenotypes. Susceptibility tests have a high measurement error; Hancock et al. (2020) estimated that the measurement error associated with the sample proportional mortality had a standard deviation (sd)=0.25 for bioassays performed using deltamethrin. Therefore, we used the predicted mean mortality to deltamethrin for *Anopheles gambiae* complex mosquitoes obtained from a series of annual predictive maps (Hancock *et al*. 2020), using the predicted value for each sample collection location and year in our analysis.

### Regression models of associations between resistance allele frequencies and mortality following exposure to deltamethrin

We assessed associations between the predicted mean mortality following exposure to deltamethrin and the predicted frequency of the *Vsgc*-995F allele. Mean mortality measurements represent the entire *Anopheles gambiae* complex, so we combined our species-specific predictions of *Vsgc*-995F frequencies across *An. gambiae, An. coluzzii*, and *An. arabiensis* to estimate the *Vsgc*-995F frequency in the *An. gambiae* complex for each sample collection location and year, *f*_*C,i*_:

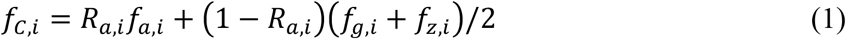

where *R*_*a,i*_ is the abundance of *An. arabiensis* at location *i* relative to the combined abundance of *An. gambiae* and *An. coluzzii*, and *f*_*a,i*_, *f*_*g,i*_ and *f*_*z,i*_ are the predicted frequencies of the *Vsgc*-995F allele in *An. arabiensis, An. gambiae*, and *An. coluzzii* at location *i*, respectively. Values of the relative abundance of *An. arabiensis* at each geographic location were obtained from the maps developed by (Sinka *et al*. 2016). We do not have spatially explicit estimates of the relative abundances of *An. gambiae* and *An. coluzzii* so we used the mean frequency across these two species in our calculation. We excluded Kenya from our regression analysis because frequencies of *Vsgc*-995F are low at our sampled locations (observed *Vsgc*-995F frequencies are less than 0.07 across 90% of samples). We tested the accuracy of our estimated *Vgsc*-995F frequencies for the *An. gambiae* complex (eq 1) against 797 of the observed *Vgsc*-995F sample frequencies in our data set that were representative of the *An. gambiae* complex (Moyes *et al*. 2019a) and found a good level of accuracy (Figure S12).

We fitted OLS linear regression models to predict mean mortality to deltamethrin using *f*_*C,i*_ as a covariate. Before model fitting, we applied the empirical logit transformation to both the independent variable and the covariate. To allow for spatial autocorrelation in the data we performed inference on the regression coefficients using cluster-robust standard errors (Conley 1999; Cameron & Miller 2015), setting the cluster radius to 100 km (Hancock *et al*. 2018).

### Importance of potential explanatory variables

In order to identify which of our potential predictor variables were having the most impact on our modelled *Vgsc* allele frequencies, we calculated measures of the importance of each predictor variable for each of the machine-learning models used in our model ensemble. It is important to note that variable importance measures cannot be used to infer causality, and they can be difficult to interpret when predictor variables are correlated. For XGB, we used the gain measure calculated for each variable using the xgboost package (Chen & Guestrin 2016), which is the fractional total reduction in the training error gained across all of that variable’s splits. For RF, we use the Gini importance, which is calculated using the sklearn package (Pedregosa *et al*. 2011). The Gini importance measures the influence of a variable in discriminating between classes in a classification algorithm (Breiman 2001). For NN, we use the permutation importance, again calculated using the sklearn package. The permutation importance of a variable is obtained by randomly shuffling the values of the variable across all observations and recalculating the model score, which in our case is the prediction error across all data points.

### Independent conditional expectation (ICE) analysis across varying ITN coverage

We studied how variation in ITN coverage impacted our model-predicted resistance allele frequencies using ICE analysis. For a single chosen location in each country, we calculated the ICE (Goldstein *et al*. 2015) of the model predicted *Vsgc*-995F frequency with varying ITN coverage for the year 2005. The ICE simply calculates the predicted response value from the model across a range of a focal predictor variable, keeping all other predictor variables fixed at their original values. This can be used to explore how the focal covariate influences the model predictions, by examining the shape and magnitude of the relationship. It is important to be aware, however, that the variation in the focal covariate is artificial and does not represent the actual variation in that particular covariate over space or time. Our ICE calculations represent variation in the model predictions for a single location and year only. The selected location within each country was chosen at random from the *Vgsc* allele frequency sampling locations for that country (the coordinates of each location are shown in Table S4). We used our model ensemble to calculated predicted *Vsgc*-995F frequencies across values of the ITN coverage in the year 2005 from zero to one in intervals of 0.1.

## ACKNOWLEDGEMENTS

We are grateful to Chantal Hendriks and Harry Gibson for their help in interpreting the predictor variables that informed our models. We thank Joseph Chabi, Edi Constant, Samuel Dadzie, Luc Djogbenou, Alex Egyir Yawson, Xavier Grau Bove, Seth Irish, Bilali Kabula, Eric Lucas, Daniel McDermott, Sanje Nagi, Eric Ochomo and Sean Tomlinson for discussions that helped in developing this work.

## Notes

### Competing Interest Statement

The authors have declared no competing interest.

