## Supplementary Figures for "Modelling spatiotemporal trends in the frequency of genetic mutations conferring insecticide target-site resistance in African malaria vector species"

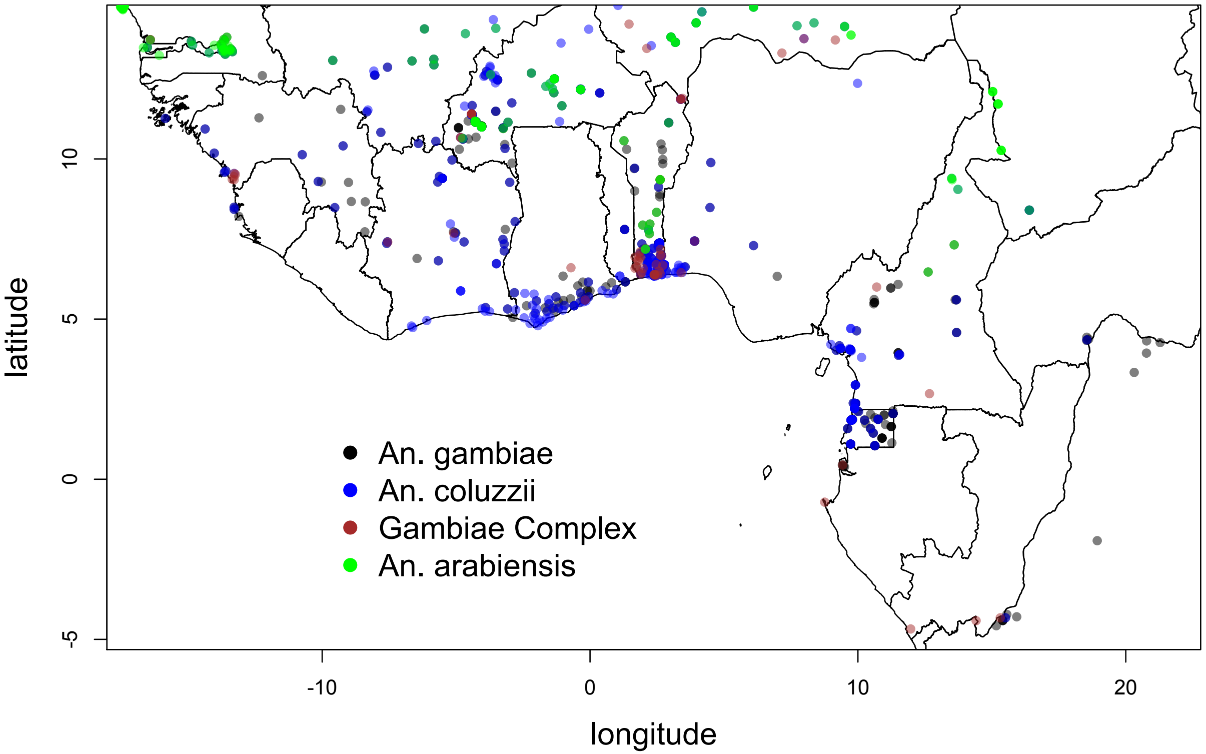

**Figure S1.** Sampling locations of the observed frequencies of the *Vgsc*-995F and *Vgsc*-995S markers in the western region of Africa that were included in our modelling analysis.

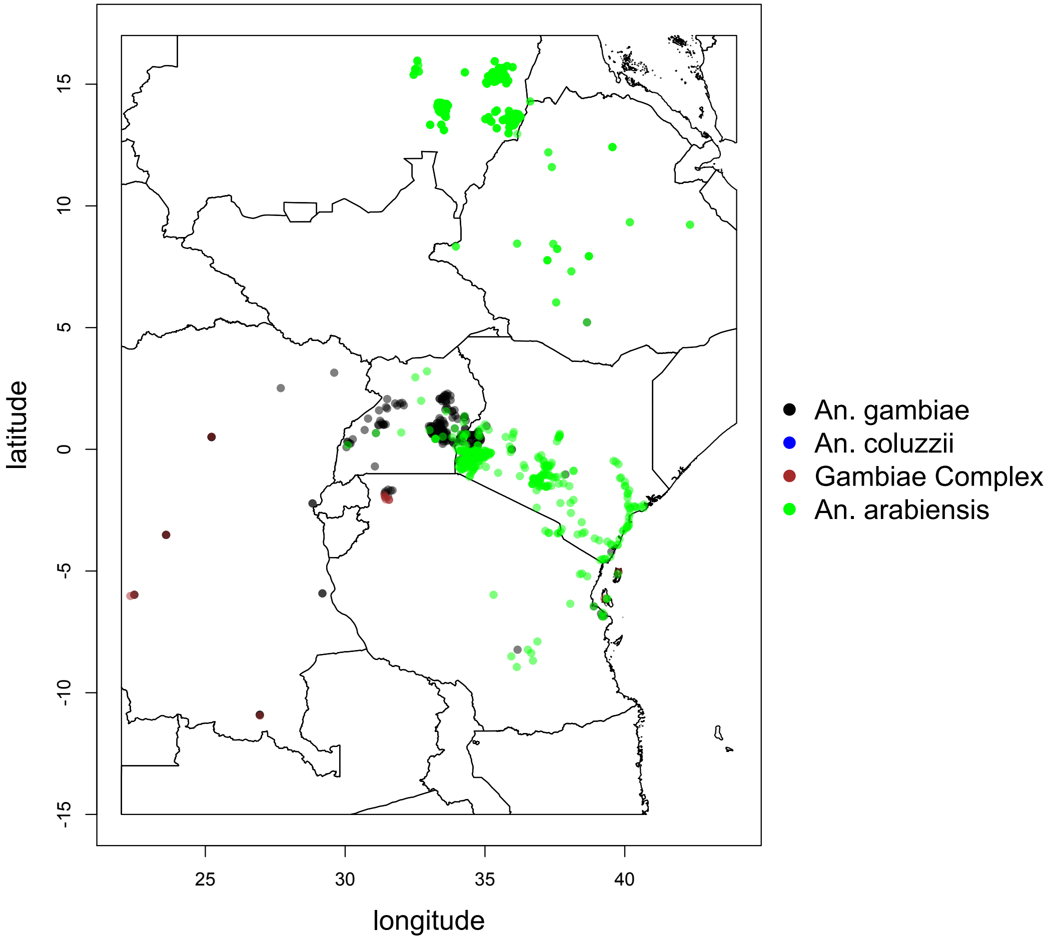

**Figure S2.** Sampling locations of the observed frequencies of the *Vgsc*-995F and *Vgsc*-995S markers in the eastern region of Africa that were included in our modellng analysis.

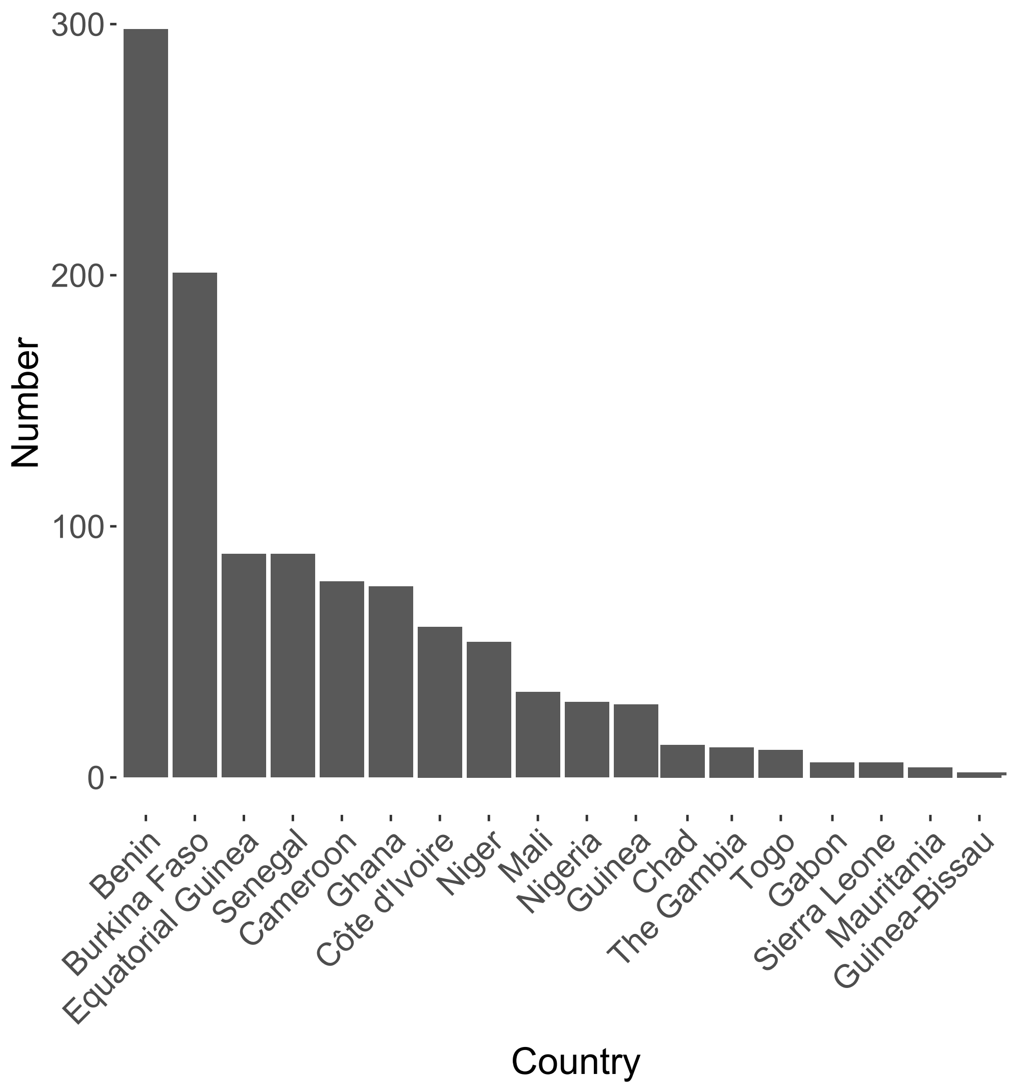

**Figure S3.** The number of samples of the frequencies of the *Vgsc*-995F and *Vgsc*-995S markers included in our data set for each country across west Africa.

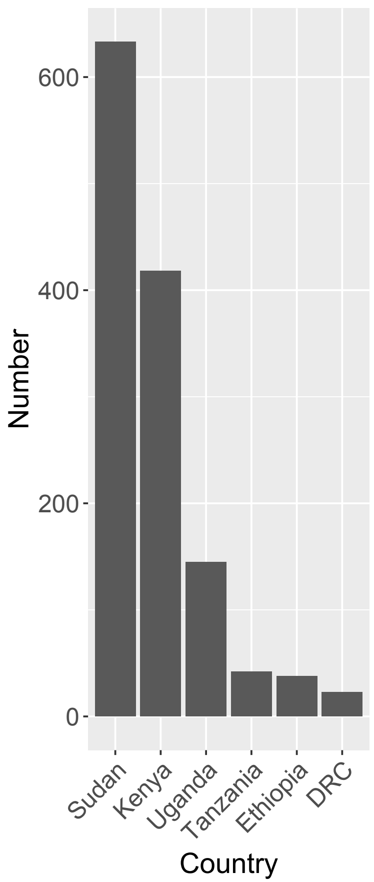

**Figure S4.** The number of samples of the frequencies of the *Vgsc*-995F and *Vgsc*-995S markers included in our data set for each country across east Africa.

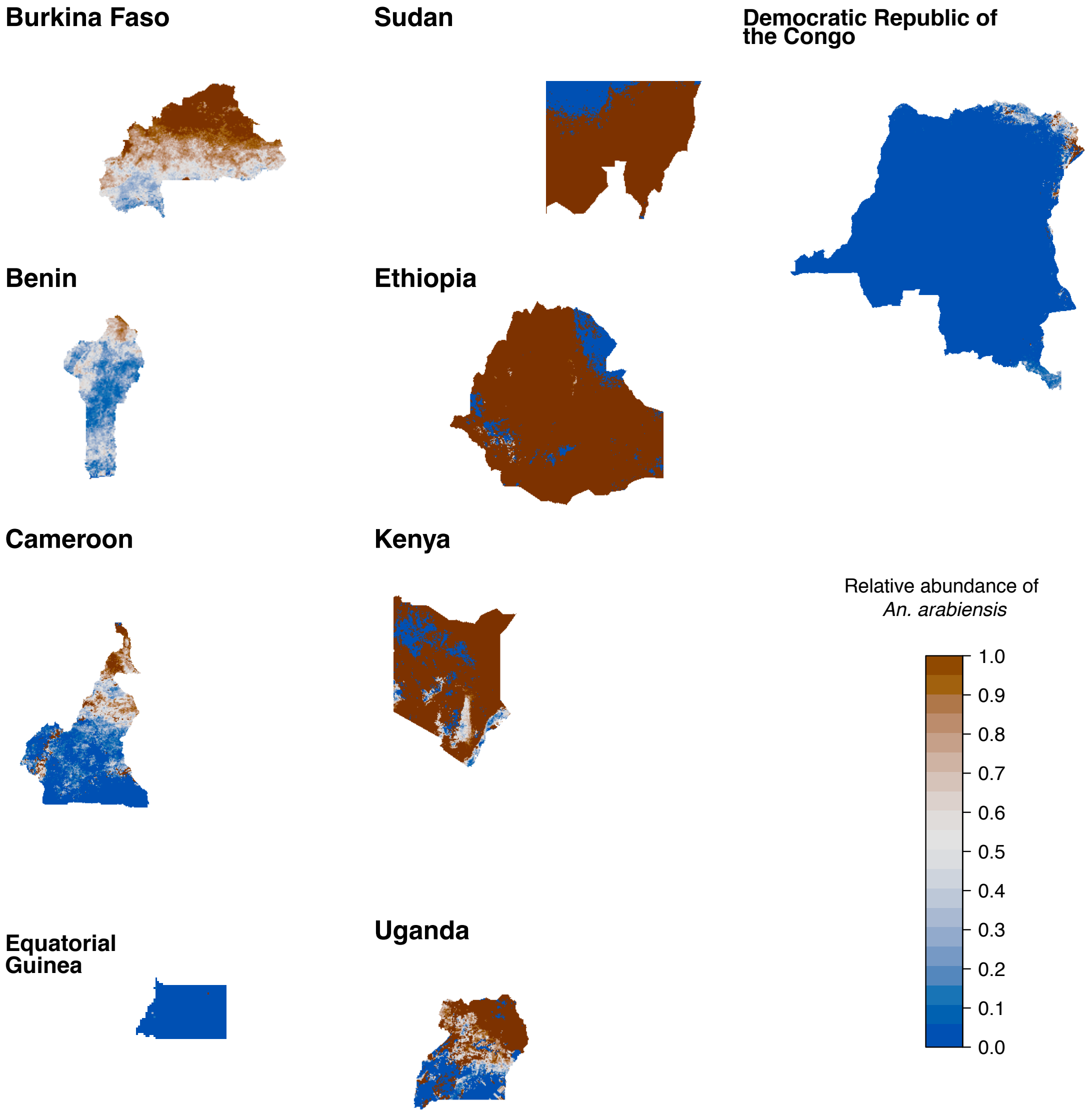

**Figure S5.** The abundance of *An. arabiensis* relative to the combined abundance of *An. gambiae* and *An. coluzzii* in the nine mapped countries. Western countries are shown in the first column from the left, eastern countries are shown in the second column from the left, and central African countries are shown in the third column from the left.

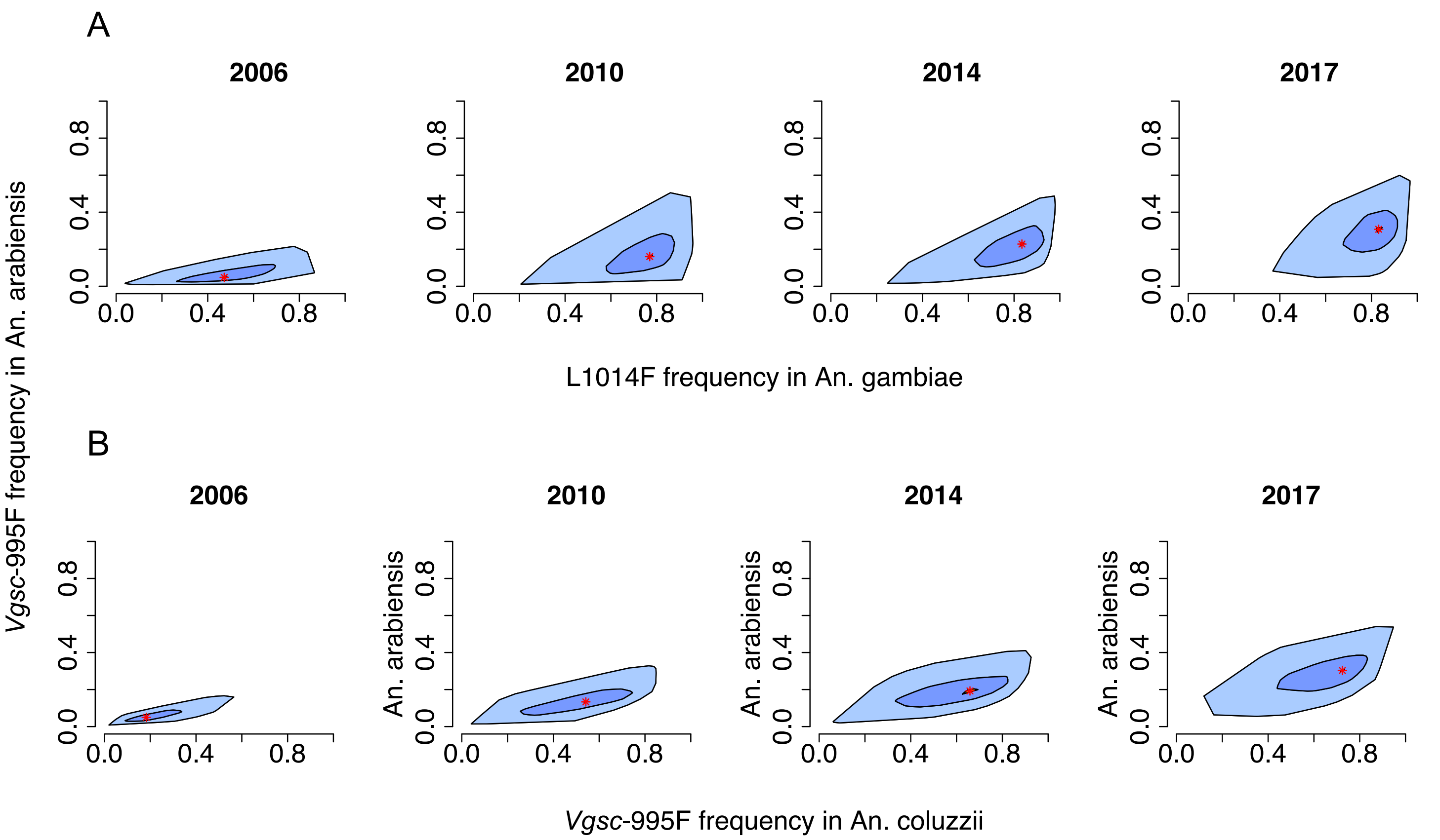

**Figure S6.** Associations between the predicted frequency of the *Vgsc*-995F allele in **A.** *An. gambiae* and *An. arabiensis*; **B.** *An. coluzzii* and *An. arabiensis.* Bagplots show the distribution across all mapped pixels within four countries in west Africa: Burkina Faso, Benin, Cameroon and Equatorial Guinea. The red asterisk shows the median, the dark blue shaded area contains 50% of all data points and the line blue shaded area contains all data points. Plots for four years are shown (from left to right): 2006, 2010, 2014 and 2017. The Pearson correlation coefficient between these predicted Vgsc-995F frequencies in *An. gambiae* and *An. arabiensis* for the years 2006, 2010, 2014 and 2017 are *r*=0.77 (CI=0.76,0.78), *r*=0.59 (CI=0.58,0.6), *r*=0.73 (CI=0.72,0.74), *r*=0.48 (CI=0.46,0.49). The Pearson correlation coefficient between these predicted *Vgsc*-995F frequencies in *An. coluzzii* and *An. arabiensis* for the years 2006, 2010, 2014 and 2017 are *r*=0.78 (CI=0.77,0.79), *r*=0.82(CI=0.81,0.83), *r*=0.74 (CI=0.73,0.75), *r*=0.66 (CI=0.65,0.67).Credible intervals were determined by boostrapping using the R package "boot".

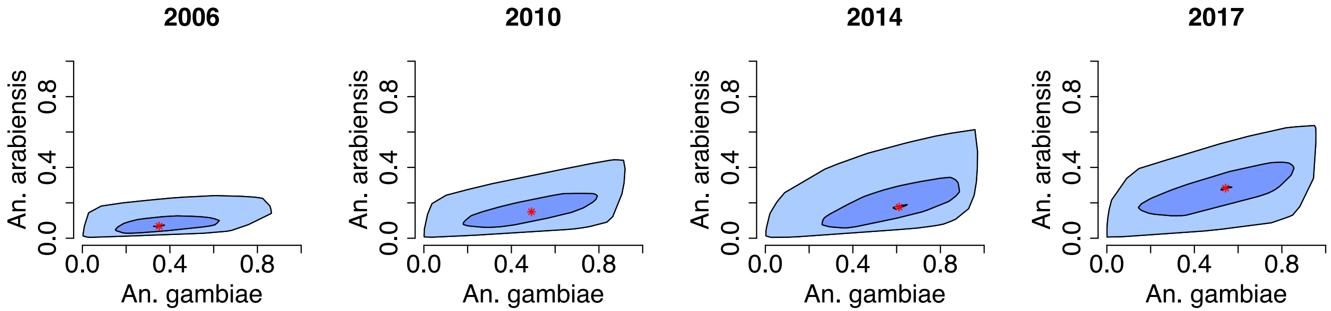

**Figure S7.** Associations between the predicted frequency of the *Vgsc*-995F allele in *An. gambiae* and *An. arabiensis.* Bagplots show the distribution across all mapped pixels within four countries in east Africa: Ethiopia, Sudan, Uganda and Kenya. The red asterisk shows the median, the dark blue shaded area contains 50% of all data points and the line blue shaded area contains all data points. Plots for four years are shown (from left to right): 2006, 2010, 2014 and 2017. The Pearson correlation coefficient between these predicted *Vgsc*-995F frequencies in *An. gambiae* and *An. arabiensis* for the years 2006, 2010, 2014 and 2017 are *r*=0.69 (CI=0.68,0.69), *r*=0.69 (CI=0.68,0.69), *r*=0.72 (CI=0.71,0.72), *r*=0.59 (CI=0.58,0.6). Credible intervals were determined by boostrapping using the R package "boot".

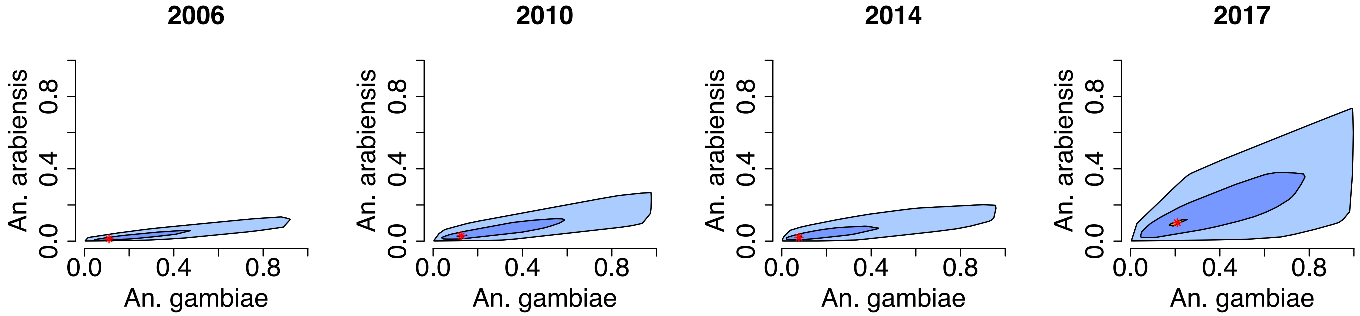

**Figure S8.** Associations between the predicted frequency of the *Vgsc*-995S allele in *An. gambiae* and *An. arabiensis.* Bagplots show the distribution across all mapped pixels within four countries in east Africa: Ethiopia, Sudan, Uganda and Kenya. The red asterisk shows the median, the dark blue shaded area contains 50% of all data points and the line blue shaded area contains all data points. Plots for four years are shown (from left to right): 2006, 2010, 2014 and 2017. The Pearson correlation coefficient between these predicted *Vgsc*-995F frequencies in *An. gambiae* and *An. arabiensis* for the years 2006, 2010, 2014 and 2017 are *r*=0.61 (CI=0.6,0.62), *r*=0.7 (CI=0.7,0.71), *r*=0.64 (CI=0.63,0.65), *r*=0.76 (CI=0.75,0.76). Credible intervals were determined by boostrapping using the R package "boot".

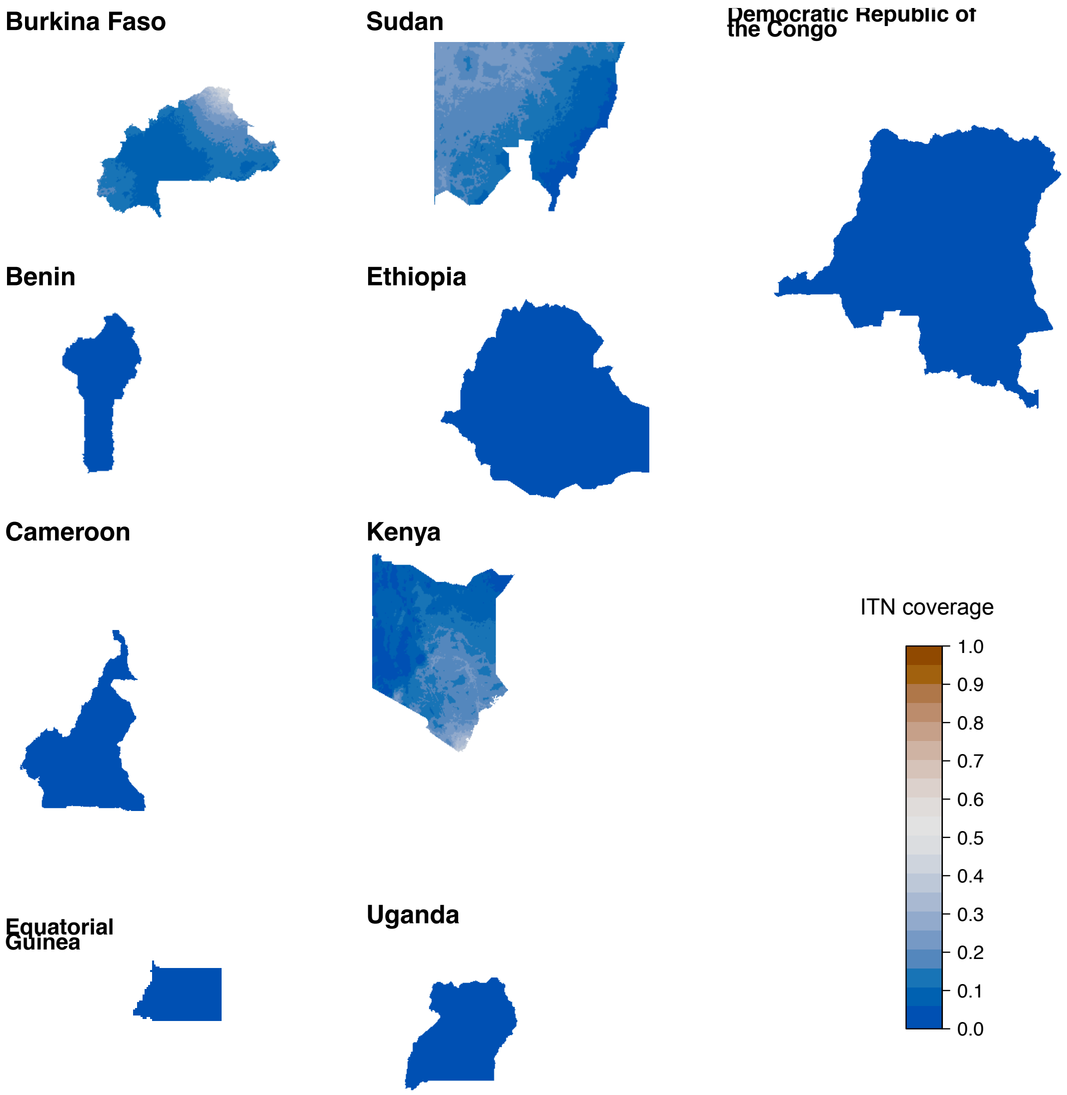

**Figure S9.** The ITN coverage in 2005 in the nine mapped countries. Western countries are shown in the first column from the left, eastern countries are shown in the second column from the left, and central African countries are shown in the third column from the left.

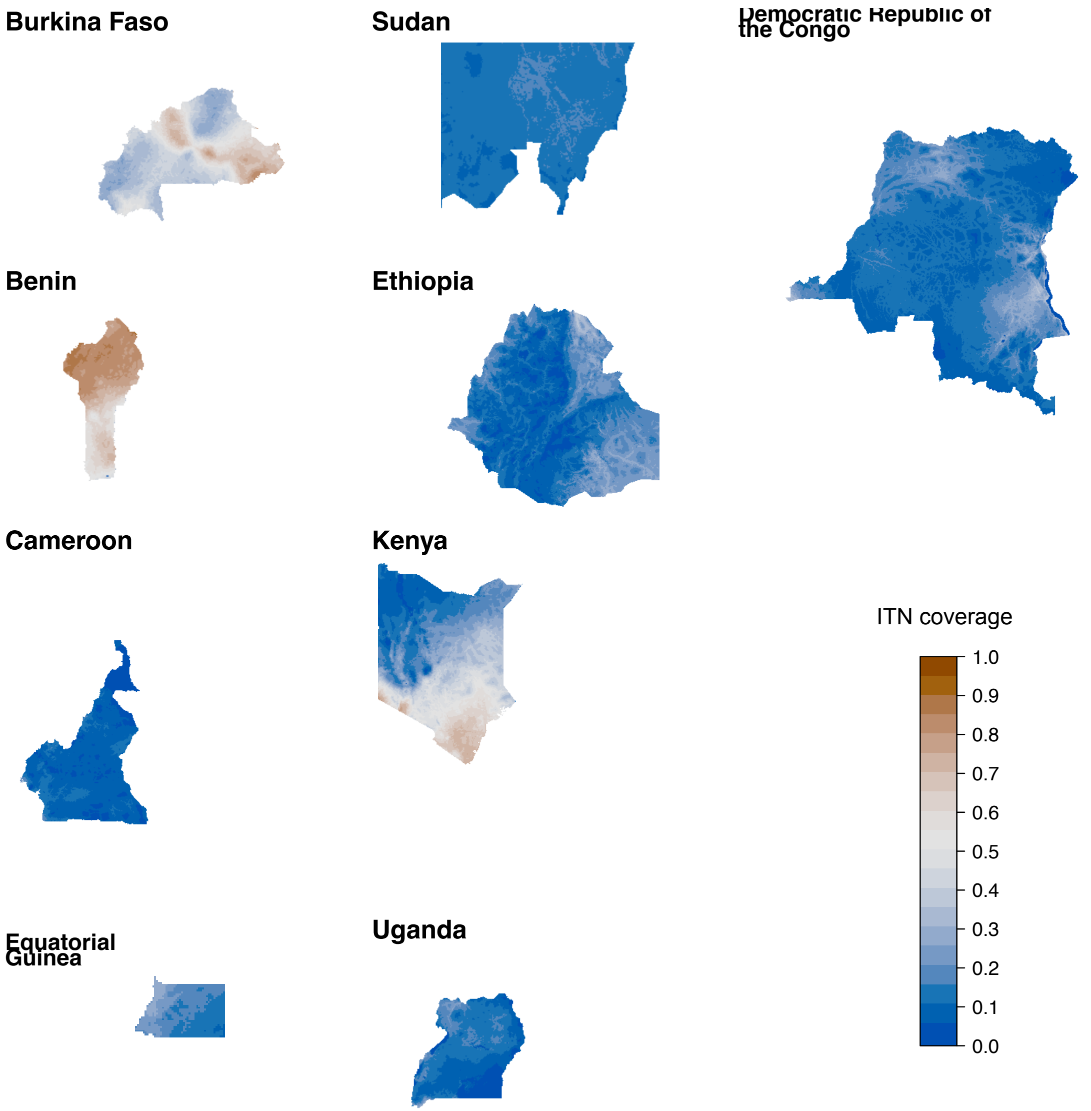

**Figure S10.** The ITN coverage in 2011 in the nine mapped countries. Western countries are shown in the first column from the left, eastern countries are shown in the second column from the left, and central African countries are shown in the third column from the left.

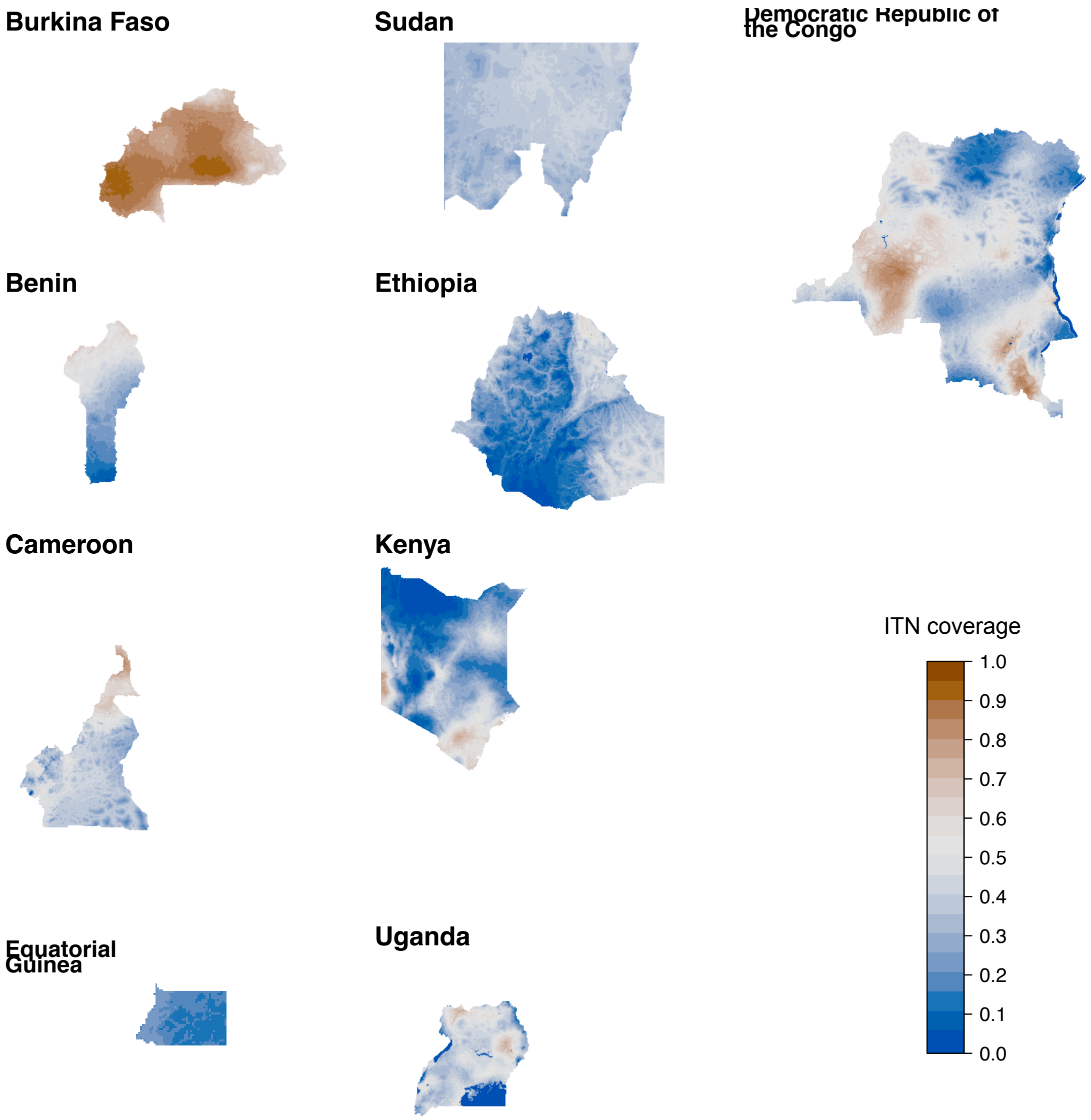

**Figure S11.** The ITN coverage in 2017 in the nine mapped countries. Western countries are shown in the first column from the left, eastern countries are shown in the second column from the left, and central African countries are shown in the third column from the left.

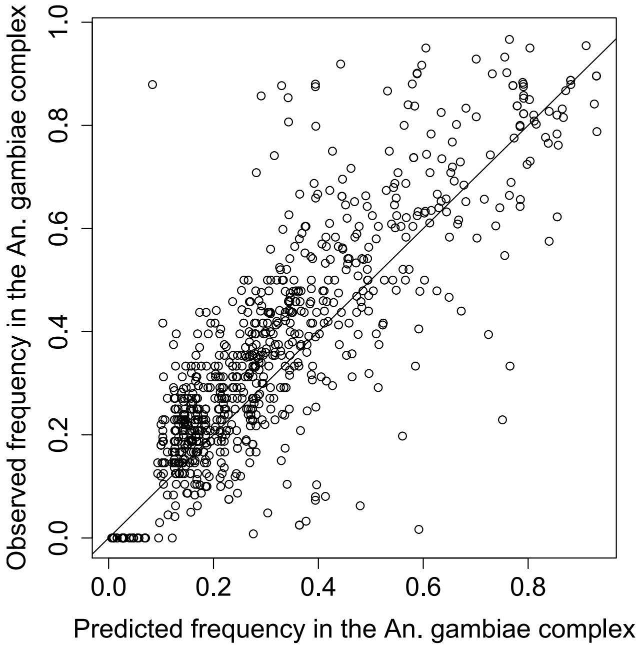

**Figure S12.** The predicted L1014F frequency in the *An. gambiae* complex derived from combining species-specific frequencies (eq #) vs observed frequencies.

**Table S1.** The root mean square error (RMSE) across the out-of-sample predictions of all *Vgsc* allele frequency observations, obtained using 10-fold cross validation. The RMSE is calculated across all observed frequencies of *Vgsc*-995L, *Vgsc*-995S and *Vgsc*-995F alleles. The out-of-sample RMSE for the multinomial logit model ensemble, together with that of each individual model constituent (XGB, RF and NN), is shown.

| **Model** | **Out-of-sample RMSE** |
|---|---|
| Multinomial logit meta-model | 0.137 |
| XGB | 0.142 |
| RF | 0.145 |
| NN | 0.155 |

**Table S2.** Countries that were included in (i) each type of mapping analysis, according to mosquito species and the type of *Vgsc* mutation mapped, and (ii) in the analysis of relationships between deltamethrin resistance phenotype and *Vgsc* mutation frequencies. For countries that were not included, the reason for exclusion is provided.

| | Mapping of ***Vgsc*-995F** frequencies in ***An. gambiae*** | Mapping of ***Vgsc*-995F** frequencies in ***An. coluzzii*** | Mapping of ***Vgsc*-995F** frequencies in ***An. arabiensis*** | Mapping of ***Vgsc*-995S** frequencies in ***An. gambiae*** | Mapping of ***Vgsc*-995S** frequencies in ***An. coluzzii*** | Mapping of ***Vgsc*-995S** frequencies in ***An. arabiensis*** | Analysis of association between mapped *Vgsc*-995F frequencies and the prevalence of mortality to deltamethrin |
|---|---|---|---|---|---|---|---|
| Burkina Faso | Yes | Yes | Yes | No. The frequency of *Vgsc*-995S is very low in west Africa (observed frequencies were zero in ~80% of the samples from the west African countries). | No. Insufficient number of samples where *Vgsc*-995S has been found in *An. coluzzii.* | No. The frequency of *Vgsc*-995S is very low in west Africa. | Yes |
| Benin | Yes | Yes | Yes | No. Same reason as for Burkina Faso. | No. Same reason as for Burkina Faso. | No. Same reason as for Burkina Faso. | Yes |
| Cameroon | Yes | Yes | Yes | No. Same reason as for Burkina Faso. | No. Same reason as for Burkina Faso. | No. Same reason as for Burkina Faso. | Yes |
| Equatorial Guinea | Yes | Yes | Yes | No. Same reason as | No. Same reason as | No. Same reason as | No. Very few deltamethrin |

| | | | | | | | |
|---|---|---|---|---|---|---|---|
| | | | | for Burkina Faso. | for Burkina Faso. | for Burkina Faso. | bioassay records are available for Equatorial Guinea. |
| Democratic Republic of the Congo | Yes | No. Insufficient number of samples representing *An. coluzzii.* | No. The relative abundance of *An. arabiensis* is very low in the DRC. | No. Insufficient number of samples measuring *Vgsc*-995S frequency. | No. Same reason as for Burkina Faso. | No. The relative abundance of *An. arabiensis* is very low in the DRC. | No. Very few deltamethrin bioassay results are available for the DRC. |
| Sudan | Yes | No. The relative abundance of *An. coluzzii* is very low in east Africa. | Yes | Yes | No. Same reason as for Burkina Faso. | Yes | Yes |
| Ethiopia | Yes | No. Same reason as for Sudan. | Yes | Yes | No. Same reason as for Burkina Faso. | Yes | Yes |
| Kenya | Yes | No. Same reason as for Sudan. | Yes | Yes | No. Same reason as for Burkina Faso. | Yes | No. The *Vgsc*-995F frequency is very low in the Kenyan samples (see Methods). |
| Uganda | Yes | No. Same reason as for Sudan. | Yes | | | Yes | No. There were insufficient deltamethrin bioassay results for Uganda. |

**Table S3.** Descriptions of each potential explanatory variable used in the ensemble model.

| Short name | Description | Temporal resolution | Lags | URL | Date accessed | Citation |
|---|---|---|---|---|---|---|
| **Insecticide-based malaria intervention coverage** | | | | | | |
| ITN coverage | ITN coverage (proportion of people protected) | annual | 0, 1, 2, 3 years | https://map.ox.ac.uk/explorer/#/ | n/a | 39 |
| ***Anopheles gambiae*** **complex species** | | | | | | |
| Arabiensis vs gambiae/coluzzi i | Proportional abundance of *An. arabiensis* to *An. coluzzii/gambiae* | static | n/a | n/a | n/a | 41 |
| **Processes associated with pesticide fate in the environment** | | | | | | |
| Leaching | Infiltration and percolation of rain or irrigation water to deeper groundwater layers. | n/a | n/a | n/a | n/a | 42 |
| Surface runoff generation | Mechanisms involved in the generation of surface runoff of rain or irrigation water. | n/a | n/a | n/a | n/a | 42 |
| Surface runoff transfer | Transfer of rain or irrigation water overland to other streams or surface water. | n/a | n/a | n/a | n/a | 42 |
| Surface runoff accumulation | Streams or surface waters where rain or irrigation water accumulates. | n/a | n/a | n/a | n/a | 42 |
| Sedimentation | Soil particles in suspension settle out of fluid, water in this instance, and come to rest. | n/a | n/a | n/a | n/a | 42 |
| Soil storage and filtering capacity | Capacity of a soil to store and filter chemical substances. | n/a | n/a | n/a | n/a | 42 |
| Volatilization† | Chemical substances convert from a liquid or solid state to a gaseous or vapour state. | monthly | n/a | n/a | n/a | 42 |
| **Crop and livestock variables** | | | | | | |
| Cropland percentage | Proportion of the pixel area covered by annual crops (temporary crops with harvest period or bare soil) | annual | 0, 1, 2, 3 years | https://modis.gsfc.nasa.gov/data/dataprod/mod12.php | 30 July 2018 | 43 |
| Cropland-natural vegetation percentage | Proportion of the pixel area covered by a mosaic of annual crops and natural vegetation (mosaic of cropland, forest, shrubland or grassland) | annual | 0, 1, 2, 3 years | https://modis.gsfc.nasa.gov/data/dataprod/mod12.php | 30 July 2018 | 43 |
| Rice | Rice production in 2005 (metric tonne) | static | n/a | http://harvestchoice.org/data/rice_p | 8 Feb 2018 | 44 |
| Cotton | Cotton production in 2005 (metric tonne) | static | n/a | https://harvestchoice.org/data/cott_p | 8 Feb 2018 | 45 |
| Sugar cane | Sugar cane production in 2005 (metric tonne) | static | n/a | https://harvestchoice.org/data/sugc_p | 8 Feb 2018 | 46 |

| | | | | | | |
|---|---|---|---|---|---|---|
| Maize | Maize production in 2005 (metric tonne) | static | n/a | https://harvestchoice.org/data/maiz_p | 8 Feb 2018 | 47 |
| Non-food | Non-food crop production in 2005 (metric tonne) | static | n/a | https://harvestchoice.org/data/area_nonf | 12 Feb 2018 | 48 |
| Banana | Banana and plantain production in 2005 (metric tonne) | static | n/a | https://harvestchoice.org/data/bapl_p | 8 Feb 2018 | 49 |
| Barley | Barley production in 2005 (metric tonne) | static | n/a | https://harvestchoice.org/data/barl_p | 8 Feb 2018 | 50 |
| Bean | Bean production in 2005 (metric tonne) | static | n/a | https://harvestchoice.org/data/bean_p | 8 Feb 2018 | 51 |
| Cassava | Cassava production in 2005 (metric tonne) | static | n/a | https://harvestchoice.org/data/cass_p | 8 Feb 2018 | 52 |
| Cereal | Cereal production in 2005 (metric tonne) | static | n/a | https://harvestchoice.org/data/cere_p | 8 Feb 2018 | 53 |
| Chickpea | Chickpea production in 2005 (metric tonne) | static | n/a | https://harvestchoice.org/data/chic_p | 8 Feb 2018 | 54 |
| Coconut | Coconut production in 2005 (metric tonne) | static | n/a | https://harvestchoice.org/data/cnut_p | 8 Feb 2018 | 56 |
| Coffee | Coffee production in 2005 (metric tonne) | static | n/a | https://harvestchoice.org/data/coff_p | 8 Feb 2018 | 57 |
| Cowpea | Cowpea production in 2005 (metric tonne) | static | n/a | https://harvestchoice.org/data/cowp_p | 8 Feb 2018 | 58 |
| Groundnut | Groundnut production in 2005 (metric tonne) | static | n/a | https://harvestchoice.org/data/grou_p | 8 Feb 2018 | 59 |
| Lentil | Lentil production in 2005 (metric tonne) | static | n/a | https://harvestchoice.org/data/lent_p | 12 Feb 2018 | 60 |
| Millet | Millet production in 2005 (metric tonne) | static | n/a | https://harvestchoice.org/data/mill_p | 12 Feb 2018 | 61 |
| Other fibres | Other fibre crop production in 2005 (metric tonne) | static | n/a | https://harvestchoice.org/data/ofib_p | 12 Feb 2018 | 63 |
| Other oils | Other oil crop production in 2005 (metric tonne) | static | n/a | https://harvestchoice.org/data/ooil_p | 12 Feb 2018 | 64 |
| Other root crops | Other roots and tubers crop production in 2005 (metric tonne) | static | n/a | https://harvestchoice.org/data/orts_p | 12 Feb 2018 | 66 |
| Palmoil | Palm oil production in 2005 (metric tonne) | static | n/a | https://harvestchoice.org/data/oilp_p | 12 Feb 2018 | 67 |
| Pigeonpea | Pigeonpea production in 2005 (metric tonne) | static | n/a | https://harvestchoice.org/data/pige_p | 12 Feb 2018 | 68 |
| Pulses | Pulses production in 2005 (metric tonne) | static | n/a | https://harvestchoice.org/data/puls_p | 12 Feb 2018 | 70 |
| Rapeseed | Rapeseed production in 2005 (metric tonne) | static | n/a | https://harvestchoice.org/data/rape_p | 12 Feb 2018 | 71 |

| | | | | | | |
|---|---|---|---|---|---|---|
| Sesame | Sesame production in 2005 (metric tonne) | static | n/a | https://harvestchoice.org/data/sesa_p | 12 Feb 2018 | 72 |
| Sorghum | Sorghum production in 2005 (metric tonne) | static | n/a | https://harvestchoice.org/data/sorg_p | 22 Apr 2018 | 73 |
| Soybean | Soybean production in 2005 (metric tonne) | static | n/a | https://harvestchoice.org/data/soyb_p | 12 Feb 2018 | 74 |
| Sunflower | Sunflower production in 2005 (metric tonne) | static | n/a | https://harvestchoice.org/data/sunf_p | 12 Feb 2018 | 75 |
| Temperate fruit | Temperate fruits production in 2005 (metric tonne) | static | n/a | https://harvestchoice.org/data/temf_p | 12 Feb 2018 | 77 |
| Tea | Tea production in 20005 (metric tonne) | static | n/a | https://harvestchoice.org/data/teas_p | 12 Feb 2018 | 78 |
| Tobacco | Tobacco production in 2005 (metric tonne) | static | n/a | https://harvestchoice.org/data/toba_p | 12 Feb 2018 | 79 |
| Tropical fruit | Tropical fruits production in 2005 (metric tonne) | static | n/a | https://harvestchoice.org/data/trof_p | 12 Feb 2018 | 80 |
| Vegetables | Vegetables production in 2005 (metric tonne) | static | n/a | https://harvestchoice.org/data/vege_p | 12 Feb 2018 | 81 |
| Wheat | Wheat production in 2005 (metric tonne) | static | n/a | https://harvestchoice.org/data/whea_p | 12 Feb 2018 | 82 |
| Yams | Yams production in 2005 (metric tonne) | static | n/a | https://harvestchoice.org/data/yams_p | 12 Feb 2018 | 83 |
| Irrigated area | Areas equipped for irrigation | static | n/a | http://www.fao.org/nr/water/aquastat/irrigationmap/index10.stm | 28 Aug 2018 | 84 |
| Harvested area | Harvested area of all crops in 2005 | static | n/a | http://harvestchoice.org/data/area_crop | 27 Feb 2018 | 85 |
| Grazing prod. | Grazing production for bovines in 2000 (kg/ha) | static | n/a | https://harvestchoice.org/data/bv_graz | 30 May 2018 | 86 |
| Stover prod. | Stover production for bovines in 2000 (kg/ha) | static | n/a | https://harvestchoice.org/data/bv_stov | 30 May 2018 | 87 |
| Cattle | Cattle density in 2005 (head/km$^2$) | static | n/a | https://harvestchoice.org/data/ad05_catt | 30 May 2018 | 88 |
| Sheep | Sheep density in 2005 (head/km$^2$) | static | n/a | https://harvestchoice.org/data/ad05_shee | 30 May 2018 | 89 |
| Goats | Goat density in 2005 (head/km$^2$) | static | n/a | https://harvestchoice.org/data/ad05_goat | 30 May 2018 | 90 |
| Pigs | Pig population in 2005 (head) | static | n/a | https://harvestchoice.org/data/an05_pig | 30 May 2018 | 91 |
| Chickens | Poultry density in 2005 (head/km$^2$) | static | n/a | https://harvestchoice.org/data/ad05_chic | 30 May 2018 | 92 |
| Livestock density | Livestock density in 2005 (LU/km$^2$) | static | n/a | https://harvestchoice.org/data/ad05_lu | 30 May 2018 | 93 |
| **Other land cover variables** | | | | | | |

| | | | | | | |
|---|---|---|---|---|---|---|
| Evergreen broadleaf percentage | Proportional cover of evergreen broadleaf forest (>60% land covered with broadleaf vegetation of height >2m and canopy never without green foliage) | annual | 0, 1, 2, 3 years | https://modis.gsfc.nasa.gov/data/dataprod/mod12.php | 30 Jul 2018 | 43 |
| Mixed forest percentage | Proportional cover of mixed forest (>60% land covered with vegetation of height >2m and mosaic of the four forest types) | annual | 0, 1, 2, 3 years | https://modis.gsfc.nasa.gov/data/dataprod/mod12.php | 30 Jul 2018 | 43 |
| Closed shrubland percentage | Proportional cover of closed shrublands (woody vegetation <2m tall with canopy cover >60% of area) | annual | 0, 1, 2, 3 years | https://modis.gsfc.nasa.gov/data/dataprod/mod12.php | 23 March 2018 | 43 |
| Open shrubland percentage | Proportional cover of open shrublands (vegetation <2m tall and shrub canopy cover >60% of area) | annual | 0, 1, 2, 3 years | https://modis.gsfc.nasa.gov/data/dataprod/mod12.php | 23 March 2018 | 43 |
| Woody savanna percentage | Proportional cover of woody savanna (trees 30-60% and understory vegetation) | annual | 0, 1, 2, 3 years | https://modis.gsfc.nasa.gov/data/dataprod/mod12.php | 23 March 2018 | 43 |
| Savanna percentage | Proportional cover of savanna (trees 10-30% and understory vegetation) | annual | 0, 1, 2, 3 years | https://modis.gsfc.nasa.gov/data/dataprod/mod12.php | 30 Jul 2018 | 43 |
| Grassland percentage | Proportional cover of grasslands (herbaceous cover with trees/shrubs <10%) | annual | 0, 1, 2, 3 years | https://modis.gsfc.nasa.gov/data/dataprod/mod12.php | 23 March 2018 | 43 |
| Permanent wetland percentage | Proportional cover of permanent wetlands (a permanent mixture of water and vegetation over extensive areas) | annual | 0, 1, 2, 3 years | https://modis.gsfc.nasa.gov/data/dataprod/mod12.php | 23 March 2018 | 43 |
| Barren and sparsely populated area percentage | Proportional cover of barren and sparsely populated areas (land with exposed soil, sand or rocks, with <10% vegetation cover at any time) | annual | 0, 1, 2, 3 years | https://modis.gsfc.nasa.gov/data/dataprod/mod12.php | 2 Aug 2018 | 43 |
| **Other variables** | | | | | | |
| Population density | Human population size (No. persons/pixel) | annual | 0, 1, 2, 3 years | https://www.worldpop.org/geodata/listing?id=17 | n/a | 94 |
| Drainage class | Classification for the rate at which water infiltrates into the soil. | n/a | n/a | http://data2.isric.org/geonetwork/srv/api/records/953d0964-6746-489a-a8d1-f188595516a9 | 9 Nov 2018 | 95 |
| Soil moisture | Moisture content of a soil (%). | n/a | n/a | https://smap.jpl.nasa.gov/data/ | 1 Dec 2018 | 96 |
| Bedrock | Depth at which bedrock occurs (cm) | n/a | n/a | https://files.isric.org/soilgrids/data/recent/ | 1 Dec 2018 | 97 |
| Flow accumulation | Based on the digital elevation model a map on flow accumulation was created. | n/a | n/a | https://hydrosheds.org/ | 26 July 2018 | 98 |
| Slope | Slope of the land (°) | n/a | n/a | https://cgiarcsi.community/data/srtm-90m-digital-elevation-database-v4-1/ | 23 March 2018 | 99 |
| Soil depth | Depth of the soil layer (cm) | n/a | n/a | n/a | n/a | 100 |
| Rainfall erosivity factor | Factor that indicates the kinetic energy of raindrop's impact and the rate of associated runoff. | n/a | n/a | https://esdac.jrc.ec.europa.eu/content/global-rainfall-erosivity | 4 Sep 2018 | 101 |
| Slope-length factor | Factor that describes the effect of slope steepness and the impact of slope length. | n/a | n/a | n/a | n/a | 42 |

| | | | | | | |
|---|---|---|---|---|---|---|
| Erosion | Total detachment and removal of soil material by water (t/ha/yr). | n/a | n/a | n/a | n/a | 42 |
| Cation exchange capacity | Cation exchange capacity of a soil is a measure for the amount of cations that can retain on soil particle surfaces ($cmol_c$/kg) | n/a | n/a | https://files.isric.org/soilgrids/data/recent/ | 21 Feb 2018 | 97 |
| Clay content | Percent of clay particles (<2μm) in the soil (%). | n/a | n/a | https://files.isric.org/soilgrids/data/recent/ | 21 Feb 2018 | 97 |
| Soil organic carbon | Organic carbon content in the soil (g/kg) | n/a | n/a | https://files.isric.org/soilgrids/data/recent/ | 21 Feb 2018 | 97 |
| Soil pH | Soil pH is a measure of acidity or alkalinity of a soil. | n/a | n/a | https://files.isric.org/soilgrids/data/recent/ | 6 Feb 2018 | 97 |
| GUF | Binary map of urban areas in 2011 (areas featuring man-made building structures with a vertical component) | static | no | https://www.dlr.de/eoc/en/desktop default.aspx/tabid-9628/16557_read-40454/ | 7 Feb 2017 | 102 |
| **Climatic variables** | | | | | | |
| Solar rad. † | Solar radiation (kJ/m²/day) | monthly | n/a | http://worldclim.org/version2 | 4 Sep 2018 | 103 |
| Wind speed† | Long-term (1970-2000) average wind speed (m/s) | monthly | n/a | http://worldclim.org/version2 | 16 Jan 2018 | 103 |
| Relative humidity | Average relative humidity (ratio of the partial pressure of water vapour to the equilibrium vapour pressure of water) between 2015 and 2018 (%) | static | n/a | https://developers.google.com/earth-engine/datasets/catalog/NOAA_GFS0P25 | 3 Dec 2018 | 104 |
| Vegetation index max† | Maximum enhanced vegetation index is a measure of greenness reflectance of the land surface | annual monthly | 0, 1, 2, 3 years | https://lpdaac.usgs.gov/products/mcd43d6*2-4*v006/ | 17 Sep 2018 | 105 |
| Vegetation index mean† | Mean enhanced vegetation index is a measure of greenness reflectance of the land surface | annual monthly | 0, 1, 2, 3 years | https://lpdaac.usgs.gov/products/mcd43d6*2-4*v006/ | 17 Sep 2018 | 105 |
| Vegetation index min† | Minimum enhanced vegetation index is a measure of greenness reflectance of the land surface | annual monthly | 0, 1, 2, 3 years | https://lpdaac.usgs.gov/products/mcd43d6*2-4*v006/ | 17 Sep 2018 | 105 |
| Land surface temp. day max† | Maximum land surface daytime temperature (°C) gap-filled from the source. | annual monthly | 0, 1, 2, 3 years | https://lpdaac.usgs.gov/products/mod11a2v006/ | 9 Oct 2018 | 106 |
| Land surface temp. day mean† | Mean land surface daytime temperature (°C) gap-filled from the source. | annual monthly | 0, 1, 2, 3 years | https://lpdaac.usgs.gov/products/mod11a2v006/ | 9 Oct 2018 | 106 |
| Land surface temp. day min† | Minimum land surface daytime temperature (°C) gap-filled from the source. | annual monthly | 0, 1, 2, 3 years | https://lpdaac.usgs.gov/products/mod11a2v006/ | 9 Oct 2018 | 106 |
| Land surface temp. diurnal diff max† | Maximum difference between corresponding surface daytime temperature and surface night-time temperature images (°C) | annual monthly | 0, 1, 2, 3 years | https://lpdaac.usgs.gov/products/mod11a2v006/ | 4 Oct 2018 | 106 |
| Land surface temp. diurnal diff mean† | Mean difference between corresponding surface daytime temperature and surface night-time temperature images (°C) | annual monthly | 0, 1, 2, 3 years | https://lpdaac.usgs.gov/products/mod11a2v006/ | 4 Oct 2018 | 106 |
| Land surface temp. diurnal diff min† | Minimum difference between corresponding surface daytime temperature and surface night-time temperature images (°C) | annual monthly | 0, 1, 2, 3 years | https://lpdaac.usgs.gov/products/mod11a2v006/ | 4 Oct 2018 | 106 |

| | | | | | | |
|---|---|---|---|---|---|---|
| Land surface temp. night max† | Maximum land surface night-time temperature (°C) | annual monthly | 0, 1, 2, 3 years | https://lpdaac.usgs.gov/products/mod11a2v006/ | 5 Oct 2018 | 106 |
| Land surface temp. night mean† | Mean land surface night-time temperature (°C) | annual monthly | 0, 1, 2, 3 years | https://lpdaac.usgs.gov/products/mod11a2v006/ | 5 Oct 2018 | 106 |
| Land surface temp. night min† | Minimum land surface night-time temperature (°C) | annual monthly | 0, 1, 2, 3 years | https://lpdaac.usgs.gov/products/mod11a2v006/ | 5 Oct 2018 | 106 |
| Rainfall† | Total precipitation (mm) | annual monthly | 0, 1, 2, 3 years | http://chg.geog.ucsb.edu/data/chirps/#_Data | 27 Nov 2017 | 107 |
| Rainfall Intensity† | Average precipitation intensity (total precipitation/No. precipitation days) (mm) | annual monthly | 0, 1, 2, 3 years | http://chg.geog.ucsb.edu/data/chirps/#_Data | 11 Dec 2018 | 107 |
| Bare surface moisture max† | Maximum values for a measure of moisture on bare surfaces (TCB, variation in soil background reflectance) | annual monthly | 0, 1, 2, 3 years | https://lpdaac.usgs.gov/products/mcd43d6*2-4*v006/ | 6 Dec 2018 | 108 |
| Bare surface moisture mean† | Mean values for a measure of moisture on bare surfaces (TCB, variation in soil background reflectance) | annual monthly | 0, 1, 2, 3 years | https://lpdaac.usgs.gov/products/mcd43d6*2-4*v006/ | 6 Dec 2018 | 108 |
| Bare surface moisture min† | Minimum values for a measure of moisture on bare surfaces (TCB, variation in soil background reflectance) | annual monthly | 0, 1, 2, 3 years | https://lpdaac.usgs.gov/products/mcd43d6*2-4*v006/ | 6 Dec 2018 | 108 |
| Surface wetness max† | Maximum values for a measure of surface moisture (TCW, variation in the vigour of green vegetation) | annual monthly | 0, 1, 2, 3 years | https://lpdaac.usgs.gov/products/mcd43d6*2-4*v006/ | 3 Oct 2018 | 108 |
| Surface wetness mean† | Mean values for a measure of surface moisture (TCW, variation in the vigour of green vegetation) | annual monthly | 0, 1, 2, 3 years | https://lpdaac.usgs.gov/products/mcd43d6*2-4*v006/ | 3 Oct 2018 | 108 |
| Surface wetness min† | Minimum values for a measure of surface moisture (TCW, variation in the vigour of green vegetation) | annual monthly | 0, 1, 2, 3 years | https://lpdaac.usgs.gov/products/mcd43d6*2-4*v006/ | 3 Oct 2018 | 108 |
| Potential evapotranspiration max | Max. potential evapotranspiration (water vapour flux under ideal conditions) between 1950 and 2000 (mm) | static | n/a | https://cgiarcsi.community/data/global-aridity-and-pet-database/ | 5 Feb 2015 | 109 |
| Potential evapotranspiration mean | Mean potential evapotranspiration (water vapour flux under ideal conditions) between 1950 and 2000 (mm) | static | n/a | https://cgiarcsi.community/data/global-aridity-and-pet-database/ | 5 Feb 2015 | 109 |
| Potential evapotranspiration min | Min. potential evapotranspiration (water vapour flux under ideal conditions) between 1950 and 2000 (mm) | static | n/a | https://cgiarcsi.community/data/global-aridity-and-pet-database/ | 5 Feb 2015 | 109 |
| Potential evapotranspiration st.dev. | Standard deviation of the potential evapotranspiration (water vapour flux under ideal conditions) between 1950 and 2000 (mm) | static | n/a | https://cgiarcsi.community/data/global-aridity-and-pet-database/ | 5 Feb 2015 | 109 |
| Elevation | Elevation measured using the hydrologically conditioned Digital Elevation Model (m) | n/a | n/a | https://hydrosheds.org/ | 27 March 2018 | 98 |
| Distance water | Distance to water, including rivers, surface waters and oceans (m) | n/a | n/a | https://hydrosheds.org/ | 5 Nov 2015 | 98 |

† Conducted a principal component analysis on variables for each month and selected the top three principal components.

**Table S4.** The location in each country for which ICE relationships between predicted *Vgsc*-995F frequencies and ITN coverage were calculated.

| Country | Latitude (°E) | Longitude (°N) |
|---|---|---|
| Burkina Faso | 11.40145 | -4.41939 |
| Benin | 9.350000 | 2.616670 |
| Cameroon | 5.969440 | 11.227220 |
| Equatorial Guinea | 1.91847 | 10.63520 |
| Ethiopia | 8.23333 | 37.58333 |
| Sudan | 14.14377 | 33.55163 |
| Uganda | 0.85707 | 33.9208 |
| Kenya | -0.08262 | 34.77468 |
| DRC | -3.5156740 | 23.59525 |
